# Non-Gaussian Normative Modelling With Hierarchical Bayesian Regression

**DOI:** 10.1101/2022.10.05.510988

**Authors:** Augustijn A.A. de Boer, Johanna M. M. Bayer, Seyed Mostafa Kia, Saige Rutherford, Mariam Zabihi, Charlotte Fraza, Pieter Barkema, Lars T. Westlye, Ole A. Andreassen, Max Hinne, Christian F. Beckmann, Andre Marquand

## Abstract

Normative modelling is an emerging technique for parsing heterogeneity in clinical cohorts. This can be implemented in practice using hierarchical Bayesian regression, which provides an elegant probabilistic solution to handle site variation in a federated learning framework. However, applications of this method to date have employed a Gaussian assumption, which may be restrictive in some applications. We have extended the hierarchical Bayesian regression framework to flexibly model non-Gaussian data with heteroskdastic skewness and kurtosis. To this end, we employ a flexible distribution from the sinh-arcsinh (SHASH) family, and introduce a novel reparameterisation and a Markov chain Monte Carlo sampling approach to perform inference in this model. Using a large neuroimaging dataset collected at 82 different sites, we show that the results achieved with this extension are equivalent or better than a warped Bayesian linear regression baseline model on most datasets, whilst providing better control over the parameters governing the shape of distributions that approach is able to model. We also demonstrate that the attained flexibility is useful for accurately modelling highly nonlinear relationships between aging and imaging derived phenotypes, which shows that the extension is important for pushing the field of normative modelling forward. All methods described here are available in the open-source `pcntoolkit`.

**Highlights:**

- We extended the Hierarchical Bayesian Regression framework for normative modelling
- Our extension allows modelling data with heteroskedastic skewness and kurtosis
- We developed a reparameterization of the SHASH distribution, suitable for sampling
- We provide the first implementation of the SHASH distribution in a fully Bayesian framework
- Results show that the extension outperforms current methods on various measures

## 1. Introduction

Brain disorders affect millions of people worldwide, and have a large impact on the quality of life of the patient and the people surrounding them. Correct treatment in the form of medication or therapy can help reduce symptoms. However, studies to understand mechanisms or develop treatments for these disorders are usually performed in a case-control framework, which provides group-level inferences in which homogeneity within clinical cohorts and clinical groups is presumed. These studies aim to detect differences between the clinical and the control (i.e. healthy) groups and are therefore limited to providing inferences about group averages (e.g. the ‘average patient’). However, it has been argued that the individual deviations from the groups’ means contain the most important information for stratification [1, 2]. An improved understanding of the heterogeneity within clinical cohorts may ultimately lead to better diagnosis and treatment.

Normative modelling provides a framework for understanding heterogeneity within clinical cohorts [2]. A normative model aims to provide estimates of centiles of variation of a specific set of phenotypes over the full range of the clinical covariates, i.e., across the lifespan, much like growth charts in pediatric medicine. Centiles of variation of imaging derived phenotypes (IDPs) are estimated, and subjects are assigned a point somewhere along the estimated centiles. The assumption is that healthy subjects typically land in areas of higher density than non-healthy subjects. The mean or ‘average’ subject is thus also a meaningful concept in normative models, but contrary to case-control studies, the deviations from that mean are interpreted as anomaly in the brain structure or function, and are seen as valuable resources for further downstream processing like analysis of underlying mechanism of brain disorders.

In probabilistic modelling, a distinction is made between epistemic and aleatoric uncertainty [3]. The former is due to uncertainty about a parameter or phenomenon, and generally becomes smaller as more data is collected, the latter is uncertainty due to the variation that is naturally present in the measured phenomenon. In normative modelling, we aim to capture and separate these forms of uncertainty to the highest possible degree. Separating all sources of aleatoric and epistemic uncertainty allows adequately controlling for nuisance variation, and guiding further decisions by relevant (biological) variation only.

As in many other fields, data availability in neuroscience has increased dramatically in recent years, and that trend will likely continue. Medical data, however, is sometimes subject to restrictions involving sharing, due to certain permissions not being granted. There is no guarantee that all data on which a model is to be trained can be centrally located. The question of how to do normative modelling on large, distributed datasets is thus a natural one, and it comes with a number of challenges. Firstly, the method must be able to handle large amounts of data. We have seen that Gaussian processes would be viable candidates for normative modelling, if it were not for their time complexity that scales cubically with the number of input data points [2]. Second, distributed datasets inevitably come with sources of nuisance variance, since every scan site has its own protocols, every scanner has its own unique noise characteristics, and there may be unknown sources of noise present in specific sites as well. In addition, it is well known that there are sex-related differences in neurobiological phenotypes. Hirearchical Bayesian Regression (HBR) can deal with these so-called batch effects in data by assuming a shared prior over batch-specific model parameters, as we describe in Appendix B. Third, neuroimaging data are complex and provide rich information about the individual from which they were derived. These data therefore need to be treated with care. Permission to share personal data is not always available, and transportation of the data is therefore not always possible. This requires federated learning; a distributed learning approach that does not require transportation of the data, only of the inferred parameters of model [4, 5]. In this study, we employ and extend the hierarchical Bayesian regression (HBR) approach for normative modelling [6, 7, 5] on non-Gaussian IDPs. The method involves defining a generative model, and inferring the posterior distribution over the model parameters using Markov chain Monte Carlo (MCMC) sampling. By absorbing and summarizing information contained in training data in learned posteriors, HBR supports federated learning; all information required to extend or adapt the model is contained in the model, eliminating the need of data transportation when using or even extending the model.

The current implementation of HBR assumes that the residuals follow a Gaussian distribution. However, the variability in many imaging phenotypes cannot be properly described by a Gaussian distribution, as they are skewed, kurtotic, or both. The issue of non-Gaussianity in neurological phenotypes has partially been addressed by Fraza et al. in [8]. This method accounts for non-Gaussianity by applying a monotonic warping function to the data [9], which yields a closed form for the marginal likelihood that can be maximised using standard optimisation techniques [10]. Here, we utilize the same warping function—the sinh-arcsinh transformation, Appendix A—but in a fully Bayesian approach, i.e., in the HBR framework, yielding full posterior distributions for all parameters. Applying the sinh-arcsinh transformation to a standard Gaussian distribution yields the SHASH distribution [11], which is detailed in section 2.1, a flexible distribution with separate parameters roughly corresponding to skew and kurtosis. Here we apply the HBR framework to the a canonical variant of SHASH distribution and a novel reparameterisation that substantially reduces dependency between the parameters, which is crucial for more efficient sampling. We assess its performance on some known problematic imaging phenotypes, and show that the HBR framework with a SHASH likelihood is able to model phenotypes in a way that was previously impossible.

The contributions of this work are; (i) a reparameterization of SHASH distribution and proposal of an MCMC sampling approach to infern this model; (ii) a thorough analysis of the HBR method with a SHASH likelihood in comparison with a warped-Bayesian linear regression as a baseline method [8]; (iii) an extensive evaluation of these methods on a large multi-site neuroimaging dataset containing estimates of cortical thickness and subcortical volume; (iv) an extension to the existing HBR implementation in the `pcntoolkit`.

The paper is structured as follows: we first provide a theoretical background of HBR, MCMC sampling, and the SHASH distribution in section 2, where the problems of using the SHASH distribution in a sampling context will also become clear. We introduce a reparameterization that addresses these problems in section 2.1.1. A framework for concisely describing common variations of HBR models is introduced in Appendix B. Then we evaluate our approach using several experiments with a large neuroimaging dataset in section 3, where we show the proposed method performs on par or better than existing methods. Then we apply the new HBR models to an even more non-linear dataset [12], where we show a clear advantage over existing methods. A discussion of the advantages and limitations of our approach follows in section 4, and we summarize all our findings in section 5.

## 2. Methods

At its heart, normative modelling involves fitting a probabilistic regression model to predict centiles of variation in a biological or psychometric response variable as a function of a set of clinically relevant covariates. The covariates are often taken to be demographic variables such as age and sex, but this is not necessarily the case and many other mappings are possible (e.g. using cognitive scores to predict brain activity). Early reports used classical Gaussian process regression [13][2] which is appealing due to its Bayesian non-parametric nature, exact inference and flexibility in modelling non-linearity with a relatively small number of parameters. However, the exact Gaussian process approach used in these works has two main limitations, namely its extremely poor computational scaling to large numbers of data points (owing largely to the cubic computational scaling required to solve the GPR predictive equations) and for the exact formulation (i.e. having an analytical solution), it is restricted to modelling Gaussian centiles. Whilst generalisations are possible within the Gaussian process literature to address these problems, these typically have other shortcomings. For example, modelling non-Gaussian noise distributions usually requires approximations which further increases computational complexity. The result is that other approaches with better computational scaling have been proposed in the neuroimaging literature. These are reviewed in Marquand et al 2019 [14] and include quantile regression and hierarchical linear regression techniques. More recently, the field has increasingly recognised the importance of modelling non-Gaussianity in the response variables, whilst using a distributional form to increase the precision with which outer centiles can be estimated [15].There are two main approaches that have been proposed to solve this problem in the neuroimaging literature, namely likelihood warping [8] and generalised additive models for location, scale and shape (GAMLSS) [10]. While these approaches are both capable of flexibly modelling non-Gaussianity and site effects, both have potential shortcomings: GAMLSS is not probabilistic due to the heavy regularisation penalties that are applied to constrain the flexible smoothers underlying the approach. Likelihood warping is probabilistic in nature, but still does not model the uncertainty associated with shape parameters and does not offer completely flexible control over the nature of the modelled distribution. Neither approach supports fully decentralised federated learning at training time although ad-hoc solutions have been proposed to allow model parameters to be transferred to a new site at test time. The approach proposed in this paper attempts to address these shortcomings. The HBR framework is a Bayesian modelling framework that assumes a generative model over the response variables *Y* in the form of a likelihood ℒ and a prior distribution *p* for each parameter *θ*_*i*_ of the likelihood. A general model can be expressed as such:

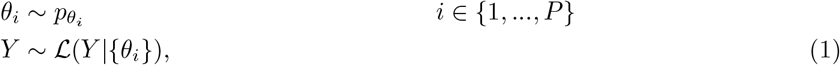

where *P* is the number of parameters of the likelihood. One of the main objectives in Bayesian modeling is to retrieve the posterior, which is the distribution over the model parameters given the data; *p*(*θ*|*Y*). Bayes’ rule gives us an expression of the posterior: *p*(*θ*|*Y*) = *p*(*Y*|*θ*)*p*(*θ*)*/p*(*Y*), but this expression can only be evaluated analytically in special cases where the likelihood is conjugate to the prior, and this is not the case in HBR. As of such, the posterior is approximated by MCMC sampling, which we discuss in section Appendix D. Assuming the response variable follows a Gaussian distribution, we substitute the Gaussian 𝒩 for ℒ, and our parameters *θ* become the mean and variance *µ* and *σ*^2^. The general model described in equation 1 is fully capable to support non-Gaussian likelihoods, like the family of SHASH distributions, which is detailed in section 2.1. The way we model *pθ*_*i*_ depends on our further assumptions about the data. Typically *µ*, and optionally also *σ* are taken to be linear functions of a set of covariates, which in this example are clinical or demographic variables. This is further detailed in Appendix B, where we also introduce a framework for model specification.

### 2.1. The SHASH Distribution

To accurately fit normative models on non-Gaussian-distributed response variables, one could substitute a family of flexible distributions for ℒ in Eq. 1. This family must then contain positively as well as negatively skewed members, leptokurtic as well as platykurtic members, and preferably with smooth transitions between them. All those requirements are fulfilled by the family of SHASH distributions (𝒮). Figures 1a and 1b show the flexibility of the 𝒮 in modelling various distributional forms.

**Figure 1:**
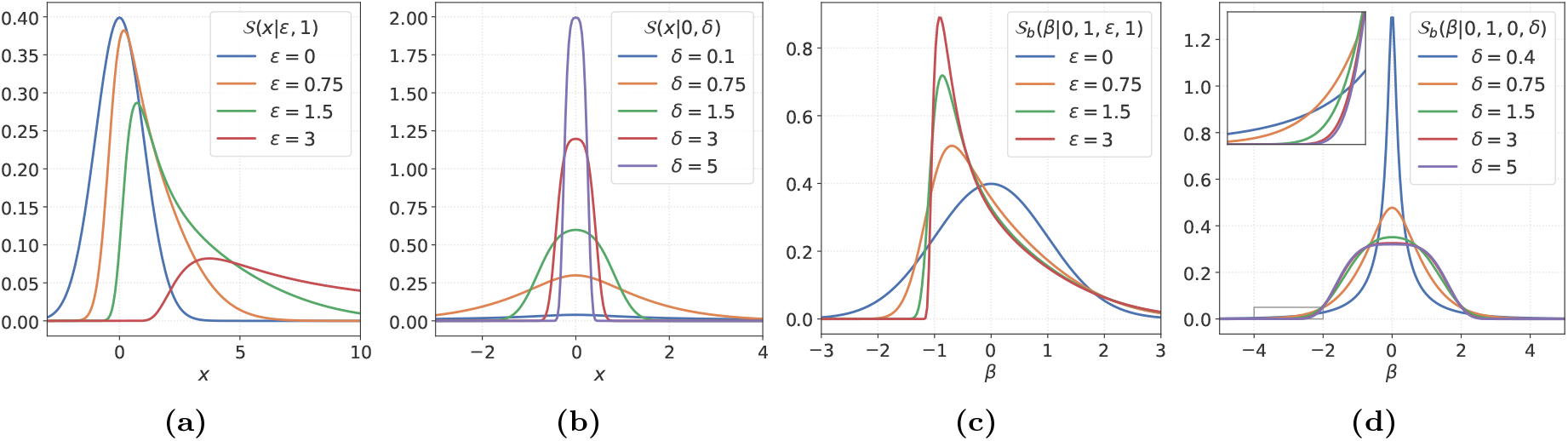
The effects of *E* and *δ* on the shape of the SHASH and SHASH_*b*_ densities. In (a) and (c), the value of *E* was varied while *δ* was kept fixed at 1. In (b) and (d), the value of *δ* was varied while *E* was kept fixed at 0. The densities in (c) and (d) have zero mean and unit variance. The inset in (d) corresponds to the area in the gray box in the bottom left corner of (d).

To generate samples from a SHASH distribution, one applies an inverse sinh-arcsinh transformation to samples from a standard Gaussian [11]. The sinh-arcsinh transformation *ξ* and its inverse *ξ*^−1^ are defined as:

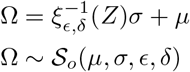

where ϵ∈ ℝ and *δ* ∈ ℝ^+^ govern the shape of the resulting distribution. Now assuming a random variable *Z* follows a standard Gaussian, we can construct samples *X* that follow a SHASH distribution by applying 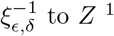:

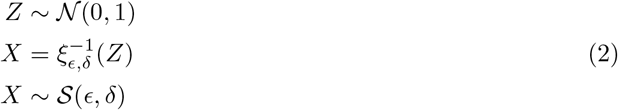

For the remainder of section 2, *Z* and *z* will indicate Gaussian distributed samples, and *X* and *x* will indicate SHASH distributed samples.

To derive the density of this SHASH distribution, we need to apply a change of variables, which states that the density of transformed samples will be the density of the original samples multiplied with the Jacobian determinant of the inverse transformation[16]. The constraint being that this Jacobian determinant exists and that it is not 0. This is true for any transformation that is a diffeomorphism^2^. Because the transformation *ξ* is a diffeomorphism, we can get the density of 𝒮 by a change of variables. We multiply the density of the samples in the Gaussian domain, *ϕ*, —which we find by applying *ξ*—with the derivative of *ξ*_*E,δ*_.

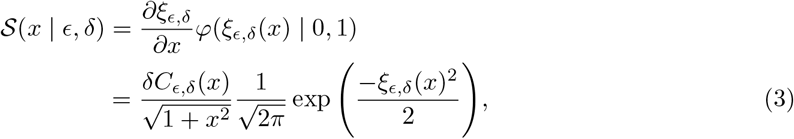

where *Cϵ*_,*δ*_(*x*) = cosh(*δ* sinh^−1^(*x*) − ϵ). Figures 1 and 2 illustrate the effect of the parameters *E* and *δ* on the shape of the SHASH density. We clearly see that *E* and *δ* modulate the skew and kurtosis, respectively.

**Figure 2:**
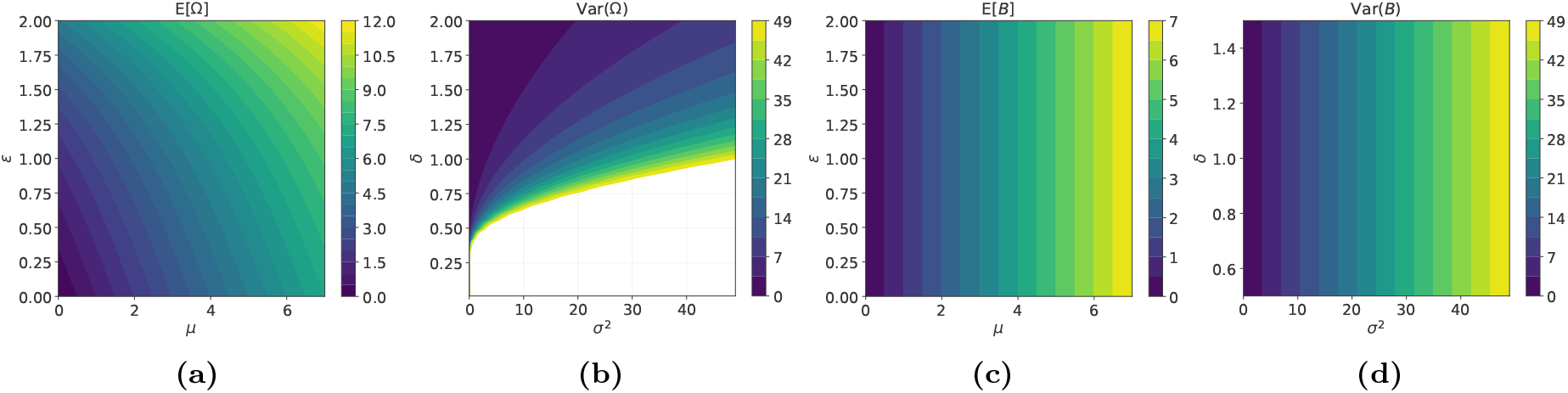
Correlations in the parameter space. In (a), at ∈= 0, the mean is exactly the value of *µ*, but when *∈ /*= 0, that is not the case. Similarly, in (b), the variance is exactly *σ*^2^ only on the line *δ* = 1. Fig. (c) and (d) show the parameter space under the proposed reparameterization. The correlation between these pairs of parameters is now removed.

A constraint on the parameter *δ* can be derived from this result. A well known property of probability distributions is that they are positive everywhere. The only term in Eq. 3 that can possibly be negative is the left fraction, because the middle fraction is a positive constant and the right exponential is strictly positive for all real inputs^3^. The left fraction is negative if exactly one of *δ* or *C*_*E,δ*_ is negative, and we have cosh(*α*) *>* 0 for all *α* ∈ ℝ. Thus the only way in which we can get a negative density is to have *δ <* 0, from which we derive the constraint that *δ >* 0.

#### 2.1.1. Reparameterization

The SHASH distribution we have seen so far has two parameters, ϵ and *δ*, but additional parameters of location and scale are necessary to achieve the desired flexibility. Jones et al. [11] suggest adding scale and location parameters by muliplication with *σ* and addition of *µ*. The resulting distribution is known as the SHASH_*o*_ or𝒮 _*o*_ distribution.

Assuming a random variable *Z* follows a standard Gaussian, we can construct samples Ω that follow a 𝒮 _*o*_ distribution by applying 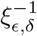 to *Z*, multiplying with *σ*, and adding *µ*:

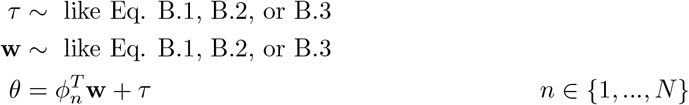

Equivalently, because of 2, we can write

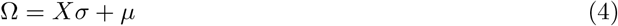

And because we already know the density of *X* to be 𝒮, we can use the chain rule in conjunction with the change of variables to arrive at the density of 𝒮_*o*_:

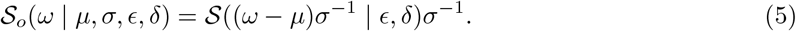

Where ϵ and *δ* are parameters that govern the higher order moments of the distribution (explained below). While we believe this family is flexible enough for the purposes of modeling a wide variety of neuroimaging phenotypes, inferring the parameters of this distribution is highly challenging because of their strong correlations^4^. Fig. 1a and Fig. 2a illustrate the correlation between *µ* and ϵ. Clearly, ϵ controls both the skew and the mean, but *µ* also controls the mean. Similarly, in Fig. 1b and Fig. 2b we see that *δ* controls the kurtosis as well as the variance, while *σ* also controls the variance. Writing down the first two moments of the 𝒮_*o*_ distribution will make this correlation explicit. Because the samples Ω from SHASH_*o*_ are simply scaled and shifted samples *X* from SHASH (see Eq. 5), we can express the moments of Ω simply in terms of the moments of *X* as:

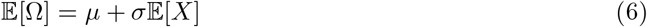

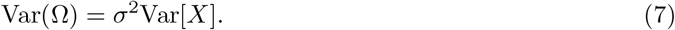

To break the correlation between *µ* and ϵ and between *σ* and *δ*, we propose to use the expressions for 𝔼 [*X*] and 𝔼 [*X*^2^] that are provided by Jones et al. to derive a new density. Jones et al. give us an analytical expression for the *r*’th non-central moment, given here in Eq. 8. For derivations of these quantities, please refer to Jones et al [11].

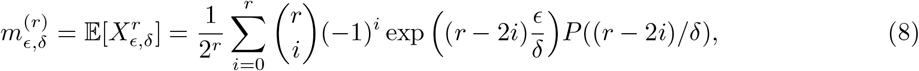

for *r* ∈ ℕ, where

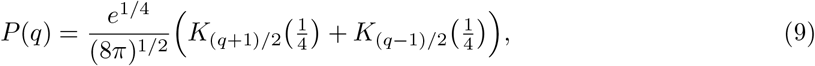

and *K* is the modified Bessel function of the second kind [17].

The solution proposed here is to first apply a standardizing shift and scale to the SHASH samples, such that the mean and variance of those samples are 0 and 1 respectively. Then, we apply the location and scale parameters *µ* and *σ* like before to attain samples *B*. Because the moments of 𝒮 are all obtainable by Eq. 8, this is a simple operation. We define the newly designed transformation as *λ* and the corresponding density as SHASH_*b*_ or 𝒮_*b*_:

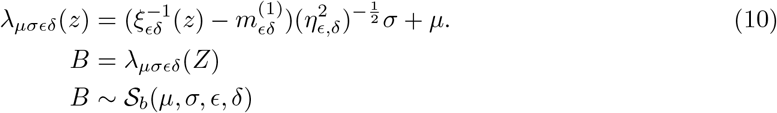

Where 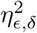 is the central variance given by 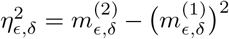. We can again use the chain rule in conjunction with the change of variables to arrive at the density of 𝒮_*b*_:

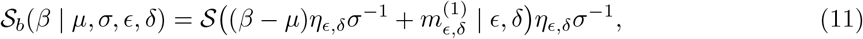

Writing down the first and second moments of *B*, the expectations that appeared in Eqs. 6 and 7 now cancel with the moments that we added in Eq. 10, and the correlation breaks completely:

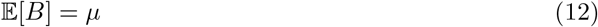

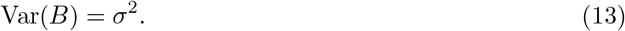

The new parameterization has nicely interpretable parameters for location and scale, which the 𝒮_*o*_ does not. Compare figures 1a and 1b with 1c and 1d. The densities in the two rightmost figures all have zero mean and unit variance. This may not seem obvious for the density with *δ* = 0.4 in Fig. 1d, but the tail behavior responsible for this becomes clearer from the inset. In Appendix G, the empirical moments of the 𝒮_*o*_ and 𝒮_*b*_ distributions are plotted over a range of parameters. It is important to note that the 𝒮_*b*_ family is isomorphic to 𝒮_*o*_, which means that they model exactly the same set of distributions, just under a different parameterization.

#### 2.1.2. Priors on 𝒮_b_ parameters

The 𝒮_*b*_ reparameterization greatly reduces the problem of correlation in parameter space, but it comes at a cost. Some of the reparameterization terms get extremely large quickly, and may therefore lead to numerical instability. This may be addressed by appropriately setting the priors for the _*b*_ likelihood. Comparing Fig. 2b and Fig. 2d, the warp in the low domain of *δ* is apparent. The stretching effect of the change of variables is enormous in those regions, because it has to compensate for the flattening effect of *δ* visible in Fig. 1b in order to standardize the SHASH distribution. The curve in the line associated with *δ* = 0.1 is barely visible in this range. The variance of *X*_*∈,δ*_ as a function of *δ* is shown on a double log scale in Fig. 3. Notice that small differences in the lower domain of *δ* amount to substantial differences in the variance. The enormous magnitude of 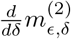 in those regions is mainly due to the behavior of the Bessel function *K*_*n*_(*z*) in Eq. 9, and to a lesser degree due to the fraction in the exponent in Eq. 8. In practical applications values of *δ <* 0.3 are thus associated with numerical instability, and sampling in those regions should be avoided at the cost of losing some expressivity. In our models we enforce this constraint by first enforcing positivity of the samples by applying the softplus function, and then adding a small constant of 0.3. The softplus function enforces positivity and is defined as:

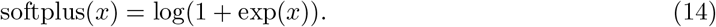

**Figure 3:**
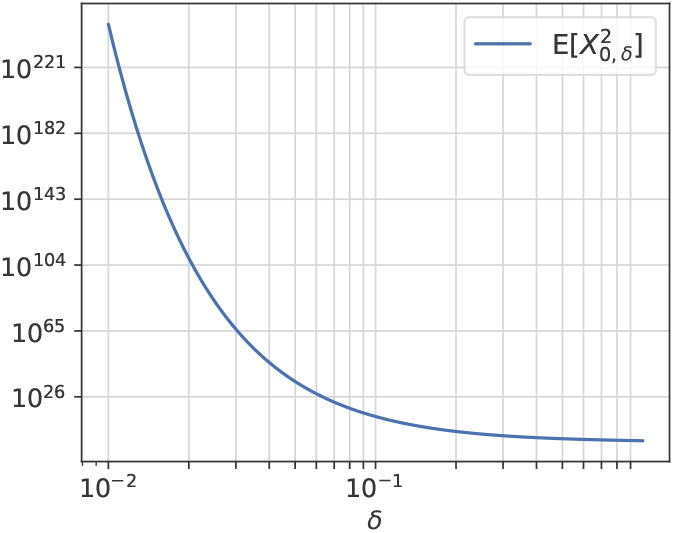
The variance of *X*_*∈,δ*_ for *δ <* 1

### 2.2. Experiments

#### 2.2.1. Dataset 1: lifespan normative modelling of image derived phenotypes

To allow a meaningful comparison with the current methods, we adapt and use the data from Rutherford et al. [18]. This is a large neuroimaging dataset collected from 82 different scanners, containing 58834 control subjects and 2925 patient subjects ^5^. Measures of cortical thickness and subcortical volume were extracted with Freesurfer version 6.0 using image derived phenotypes derived either from the Destrieux cortical parcellation or from the subcortical segmentation. Furthermore, the different UK-Biobank datasets [19] were merged, because based on prior work [8] [20], we have observed that site effects in the UKB cohort are minimal for these freesurfer measure derived from the different scanners used (due to a careful harmonisation of acquisition parameters and scanning hardware across this cohort). We therefore consider that we can safely consider them to be drawn from a single site. Counts per site and sex can be found in section Appendix F. For full details about the data, we refer to Rutherford et al. [18]. Most data in this dataset are publicly available, but for several of the clinical samples, we do not have explicit permission for data sharing. Such permissions for sharing were not obtained from the data from the University of Michigan, and as such these data were not included in this study (n=1394 (1149 controls, 245 patients)).

The family of SHASH distributions has the flexibility to model skewed and kurtotic distributions, but also has the standard Gaussian as a central member. We pick some ‘hard’ (i.e. difficult) phenotypes, the age-conditioned distributions of which are not described well by a Gaussian, and some ‘easy’ phenotypes, the age-conditioned distributions of which are described well by a Gaussian. We aim to model both easy and hard phenotypes with the same model, using the same hyperparameters. The selected hard phenotypes are: right cerebellum white matter, because it is slightly skewed; right lateral ventricle, because it is moderately skewed and heteroskedastic, and white matter hypointensities, because it is severely skewed and heteroskedastic. For easy phenotypes, we selected the estimated total intracranial volume, the thickness of the Jensen sulcus, and the volume of the brain stem. These ‘easy’ phenotypes do not show a lot of skew or kurtosis, but the mean of the thickness of the Jensen sulcus shows an interesting trajectory (see Fig. I.26). The distributions of the different phenotypes are shown in figure 6 and in section Appendix I below. In some instances we used the following abbreviations for the ‘hard’ phenotypes: Right-Cerebellum-White-Matter: RCWM, Right-Lateral-Ventricle: RLV, WM-hypointensities: WMH, and similarly for the ‘easy’ phenotypes: EstimatedTotalIntraCranialVol: ETICV, rh S interm prim-Jensen thickness: RIPJ, Brain-Stem: BS,

A stratified 10-fold split was made using the Sklearn package [21], to which end we filtered out data from sites with less than 10 subjects of one or either sex. This ensures that the test folds contain at least one unique sample for each batch effect. As preprocessing, all data was feature-wise-normalized by subtracting the mean and dividing by the standard deviation. Normalization is not a necessity, but it allowed the same hyperparameters to be used for different response variables. This would otherwise not be possible, because the scale of the response variables differ by several orders of magnitude. Sampling performance could be hampered if the prior distribution differs exceedingly much from the likelihood.

#### 2.2.2. Batch effects

Age and sex were used as covariates, and site was used as a batch effect. The HBR framework supports multidimensional batch effects, allowing to use both sex and site as a batch effect and allowing site effects to be modelled in a single step regression together with effects of interest. However, we consider that there is little reason to assume that the differences related to sex vary much between sites, and having this multidimensional batch effect results in a larger number of model parameters. In practice, we saw no difference in performance between models that used sex as a batch effect, and those that used sex as a covariate. For further discussion, see section 4.

#### 2.2.3. Dataset 2: Modelling latent representations derived from an autoencoder

As a second, more challenging, test of the flexibility of our model, we fit a normative model on a highly non-linear representation of brain data derived from the latent representation learned by a convolutional autoencoder. Full details are provided in Zabihi et al. [12], but briefly; data derived from the UK Biobank sample (n=20,781, 47% female) was compressed to a 2-dimensional latent representation using a pipeline consisting of a conditional convolutional autoencoder with a 100-dimensional latent space [22], followed by a UMAP dimensionality reduction [23]. The two-step pipeline was constructed because the intermediate representation produced by the autoencoder would otherwise not be reflective of age and sex, it would only reconstruct the data. In this case it was desirable to have a latent representation conditioned on these variables. The resulting representation followed a distribution with a nonlinear trajectory with respect to age n the mean, variance, and skew. A stratified train/test (75%/25%) split was made on the raw data level, specifically not at the 100-dimensional latent representation (see Fig. 9). This means that the same train data was used for training the convolutional autoencoder as well as the UMAP embedding, and all of the test data was held out from training. Following the rationale described above, no batch effects were modeled. Again, we used age and sex as covariates.

#### 2.2.4. MCMC Samplers

We employed a No U-Turn sampler for inference, implemented within the PyMC version 5.4.1 software using 2 chains per sample. Chains were run for 1500 iterations using 500 samples as burn-in. Convergence checks were performed using the 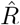 statistic. Further details about the theory behind MCMC inference are provided in Appendix D. All scripts necessary for running the analyses conducted in this paper are freely available via GitHub.

#### 2.2.5. Ethics Statement

All participants provided written informed consent for their data to be used, and all studies were approved by the institutional review boards at the respective institutions.

## 3. Results

In section 3.1, the convergence of the MCMC samplers utilized by the HBR models is assessed.

We evaluate four variants of the HBR method: The first uses a simple Gaussian likelihood, where *µ* and *σ* are modeled exactly like *µ* and *σ* in section Appendix B.3. This model is indicated with an 𝒩 . Model 𝒩 is the model that was previously the limit of what could be done within HBR in the `pcntoolkit`. We also evaluate a model with an *𝒮*_*o*_ likelihood and two variants of the *𝒮*_*b*_ likelihood, which differ only in the way *∈* and *δ* are modeled. In the first model *𝒮*_*b*1_, both *∈* and *δ* assume constant values throughout the full range of covariates. In model *𝒮*_*b*2_, both *∈* and *δ* are linear regressions on the design matrix Φ, which in our case contains 5-knot b-spline basis expansion[24] of the age covariate, concatenated with the age and sex covariates. These models are formalized in table 1, using the notation detailed in Appendix B. Details about the priors can be found in table 2. Full specifications of the generative models can be found in Appendix C.

**Table 1:** The generative descriptions of the models that are compared to W-BLR, using the notation detailed in Appendix B. For all *n, n* ∈ *{*1, …, *N }*. The softplus function was scaled down by a factor of 10 to force more linear behavior between 0 and 1. More details about the priors can be found in table 2. The notation *ℳ*_*a.b.c*_ is used to denote different generative models, the details of which can be found in Appendix C.

| $\mathcal{N}$ | $\mathcal{S}_{o1}$ | $\mathcal{S}_{b1}$ | $\mathcal{S}_{b2}$ |
| --- | --- | --- | --- |
| $\mu_n \sim \mathcal{M}_{3.1.2b}$ | $\mu_n \sim \mathcal{M}_{3.1.2b}$ | $\mu_n \sim \mathcal{M}_{3.1.2b}$ | $\mu_n \sim \mathcal{M}_{3.1.2b}$ |
| $\sigma_n^\pm \sim \mathcal{M}_{3.1.1}$ | $\sigma_n^\pm \sim \mathcal{M}_{3.1.1}$ | $\sigma_n^\pm \sim \mathcal{M}_{3.1.1}$ | $\sigma_n^\pm \sim \mathcal{M}_{3.1.1}$ |
| $\sigma_n = \text{softplus}(\sigma_n^\pm)$ | $\sigma_n = \text{softplus}(\sigma_n^\pm)$ | $\sigma_n = \text{softplus}(\sigma_n^\pm)$ | $\sigma_n = \text{softplus}(\sigma_n^\pm)$ |
| $y_n \sim \mathcal{N}(\mu_n, \sigma_n)$ | $\epsilon \sim \mathcal{M}_1$ | $\epsilon \sim \mathcal{M}_1$ | $\epsilon \sim \mathcal{M}_{3.1.1}$ |
| | $\delta^\pm \sim \mathcal{M}_1$ | $\delta^\pm \sim \mathcal{M}_1$ | $\delta^\pm \sim \mathcal{M}_{3.1.1}$ |
| | $\delta = \text{softplus}(10 \cdot \delta^\pm)/10$ | $\delta = \text{softplus}(10 \cdot \delta^\pm)/10$ | $\delta = \text{softplus}(10 \cdot \delta^\pm)/10$ |
| | $y_n \sim \mathcal{S}_o(\mu_n, \sigma_n, \epsilon, \delta + 0.3)$ | $y_n \sim \mathcal{S}_b(\mu_n, \sigma_n, \epsilon, \delta + 0.3)$ | $y_n \sim \mathcal{S}_b(\mu_n, \sigma_n, \epsilon, \delta + 0.3)$ |

**Table 2:** Prior settings for all HBR models used in this report. Note that parameters with a *±* superscript are mapped by a softplus function post-sampling to force positivity, as displayed in table 1. *𝒩* ^+^ indicates the positive Normal distribution. **0**_*D*_ and **I**_*D*_ denote a *D* dimensional zero-vector and identity matrix, respectively. Central values were chosen for most priors, with *δ* being a notable exception. As discussed before, having a low *δ* leads to numerical issues, and by setting the priors up like this, we aim to push the sampler away from the low domain. Remember that we also add a constant of 0.3 to the result of applying the softplus function to *δ*^*±*^. The prior on *τ*_*σ*_ is centered at 1, because due to standardization we expect *σ* to lay close to 1 on average, and the softplus approximates an identity function for positive values, so the mode of the transformed value should still lay close to 1. We set the prior on the weights of *µ* a little wider than the others, because we found that improved the regression. Making the priors any wider than this did not lead to any significantly different results.

| Parameter | Prior | $\mathcal{N}$ | $\mathcal{S}_{o1}$ | $\mathcal{S}_{b1}$ | $\mathcal{S}_{b2}$ |
| --- | --- | --- | --- | --- | --- |
| $\mathbf{w}_\mu$ | $\mathcal{N}(\mathbf{w}_\mu \mid \mathbf{0}_D, \mathbf{I}_D)$ | X | X | X | X |
| $\mu_{\tau_\mu}$ | $\mathcal{N}(\mu_{\tau_\mu} \mid 0, 1)$ | X | X | X | X |
| $\sigma_{\tau_\mu}$ | $\mathcal{N}^+(\sigma_{\tau_\mu} \mid 1)$ | X | X | X | X |
| $\nu_{\tau_\mu}$ | $\mathcal{N}(\nu_{\tau_\mu} \mid 0, 1)$ | X | X | X | X |
| $\mathbf{w}_{\sigma^\pm}$ | $\mathcal{N}(\mu_{\mathbf{w}_{\sigma^\pm}} \mid \mathbf{0}_D, \mathbf{I}_D)$ | X | X | X | X |
| $\tau_{\sigma^\pm}$ | $\mathcal{N}(\tau_{\sigma^\pm} \mid 1, 1)$ | X | X | X | X |
| $\epsilon$ | $\mathcal{N}(\epsilon \mid 0, 1)$ | | X | X | |
| $\mathbf{w}_\epsilon$ | $\mathcal{N}(\mathbf{w}_\epsilon \mid \mathbf{0}_D, 0.2 \cdot \mathbf{I}_D)$ | | | | X |
| $\tau_\epsilon$ | $\mathcal{N}(\tau_\epsilon \mid 0, 0.2)$ | | | | X |
| $\delta^\pm$ | $\mathcal{N}(\delta^\pm \mid 1, 1)$ | | X | X | |
| $\mathbf{w}_\delta$ | $\mathcal{N}(\mathbf{w}_\delta \mid \mathbf{0}_D, 0.2 \cdot \mathbf{I}_D)$ | | | | X |
| $\tau_\delta$ | $\mathcal{N}(\tau_\delta \mid 1, 0.3)$ | | | | X |

In section 3.2, we compare different types of HBR normative models, with one another and with warped Bayesian Linear Regression [8], which is used as a baseline, abbreviated with W-BLR. The models used in section 3.3 are the same as in section 3.2, just applied to different data.

Note that the results in section 3.2 are based on MCMC estimates of z-scores (see section Appendix E). The results in section Appendix C are based on the MAP estimates, which, were chosen because they are less computationally demanding, and because in practice, the MCMC results were only marginally better.

A main objective in Normative modelling is to map observations to z-scores in such a way that healthy variation within those z-sores follows a Gaussian distribution, and that patients fall in the outer centiles of variation. Gaussianity in the outer centiles is compared by the third and fourth moment (skew and kurtosis) of the predicted z-scores. The qq-plots, and a visual inspection of the regression curves further help assess the centiles.

### 3.1. Convergence Analysis

Here a simple analysis of the samplers is performed, by analyzing the trajectory of the Gelman-Rubin, or 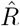 statistic [25], which can be thought of as the ratio between the interchain variance and the intrachain variance. The 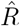 statistic is 1 (one) when those are equal, and this is a good indicator that the sampler has converged. In practice, a rule of thumb is that an 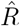 lower than 1.1 is no reason to worry, and an 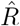 less than 1.05 indicates good convergence. To compute the 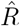 statistic, at least two chains need to be sampled. We sampled two chains of 1500 samples using the `pcntoolkit`, the first 500 of which were used for tuning and were discarded. The last 1000 samples of all chains were stored and used for further analysis.

In figure 4, the 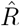 statistics of *∈* (unbroken line) and *δ* (broken line) are displayed as a function of chain length. The results are averaged over the 10 folds. Overall, the convergence of the *𝒮*_*o*1_ model, the *𝒮*_*b*1_ model and the 𝒮_*b*2_ models are similar on these phenotypes. Except for the `WM-hypointensities` phenotype - which is very hard to model - all samplers converge with 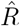 very close to 1 within at most 300 iterations.

**Figure 4:**
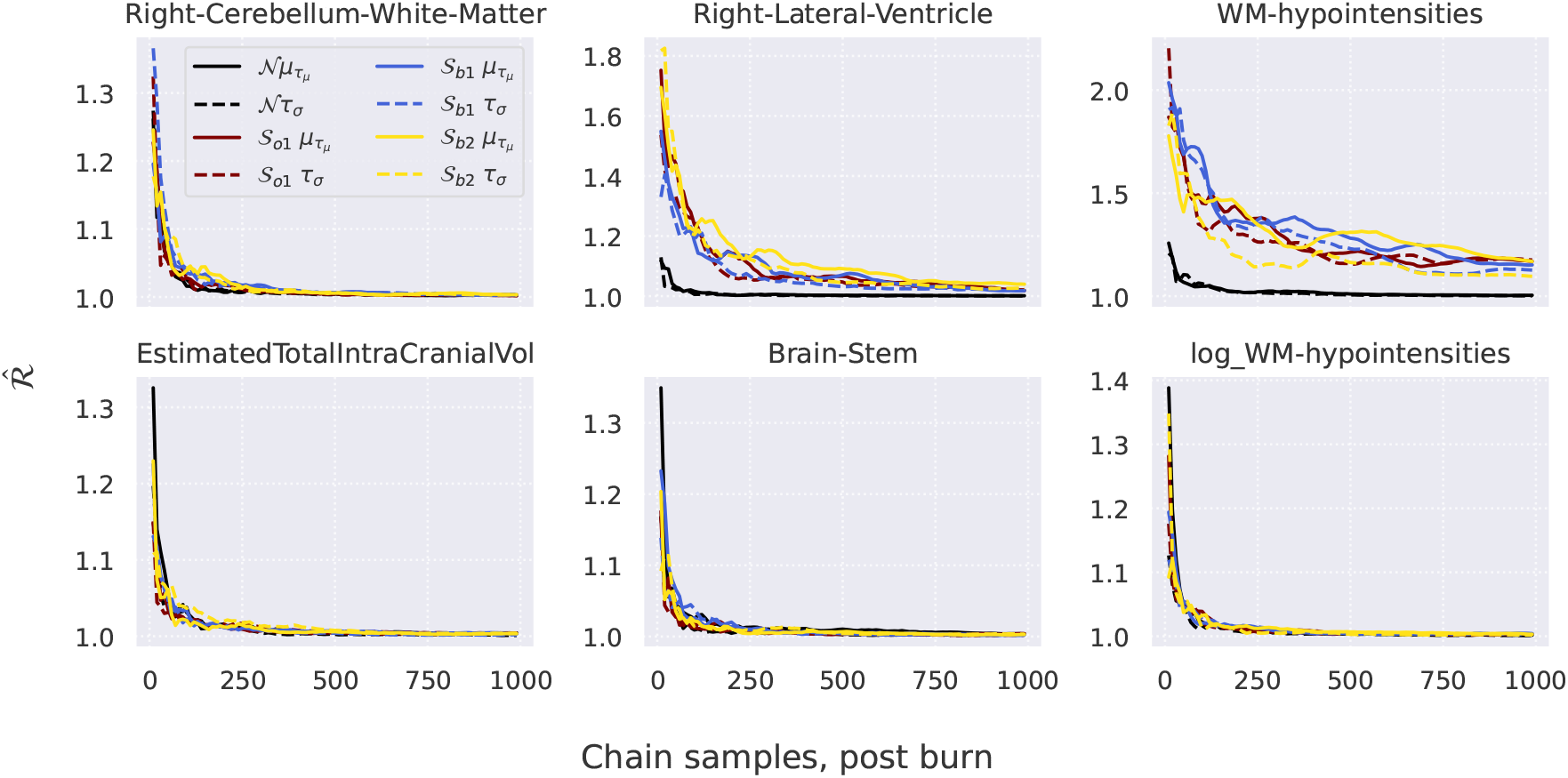
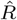 of *∈* and *δ* for *𝒮*_*o*_, *𝒮*_*b*1_ and *𝒮*_*b*2_, average of 10 folds. For the selection of ‘easy’ phenotypes in this panel, all models behave comparably. For the selection of ‘hard’ phenotypes, the *𝒮*_*o*_, *𝒮*_*b*1_, *𝒮*_*b*2_ models behave comparably. ‘Hard’ phenotypes: Right-Cerebellum-White-Matter, Right-Lateral-Ventricle, (log)White-Matter-Hypointensities. ‘Easy’ phenotyopes: Estimated-Total-Intracranial-Volume, Brain-Stem. See also section 2.2.

Since it is a strictly positive measure, we also evaluate the convergence of the samplers on a log-transformed version of the `WM-hypointensities`, which shows much better convergence.

Note also that the *𝒮*_*o*_ sampler did not converge for the most difficult phenotype reported in section 3.3, so we use the *𝒮*_*b*_ parameterisation in that case.

### 3.2. Goodness-of-fit Analysis

First, we discuss how the HBR with the *𝒮*_*b*_ likelihoods compare to the W-BLR method, and the HBR with the Gaussian likelihood. We assess (i) how well the predicted z-scores follow a Gaussian distribution, by looking at the higher-order moments of the z-scores and at qq-plots; (ii) how good the fit of the percentile lines is visually, by superimposing them on the data, and (iii), how well the models have captured the batch effects, by looking at classification performance. Specifically: for any combination of 2 sites, we computed the area-under-the-curve (AUC) metric [26] by using the z-scores as classification thresholds, and the batch-indices as labels.

Table 3 forms the first basis of our discussion. For convenience, it is useful to juxtapose figure 6 or Appendix I, which visualize the phenotypes discussed here. The skew and kurtosis are the third and fourth standardized moment respectively [27]. As the name suggests, skew describes how much the data is skewed about the mean. The kurtosis measures the tailed-ness of the distribution. A spike distribution has a kurtosis of 0, and a standard Gaussian has a kurtosis of 3. For this reason, the kurtosis values reported here are reduced by 3, which is sometimes named the *excess kurtosis*. For both measures, the ideal result is thus 0, which suggests a symmetric distribution, with tails as slim as those of a Gaussian.

**Table 3:**
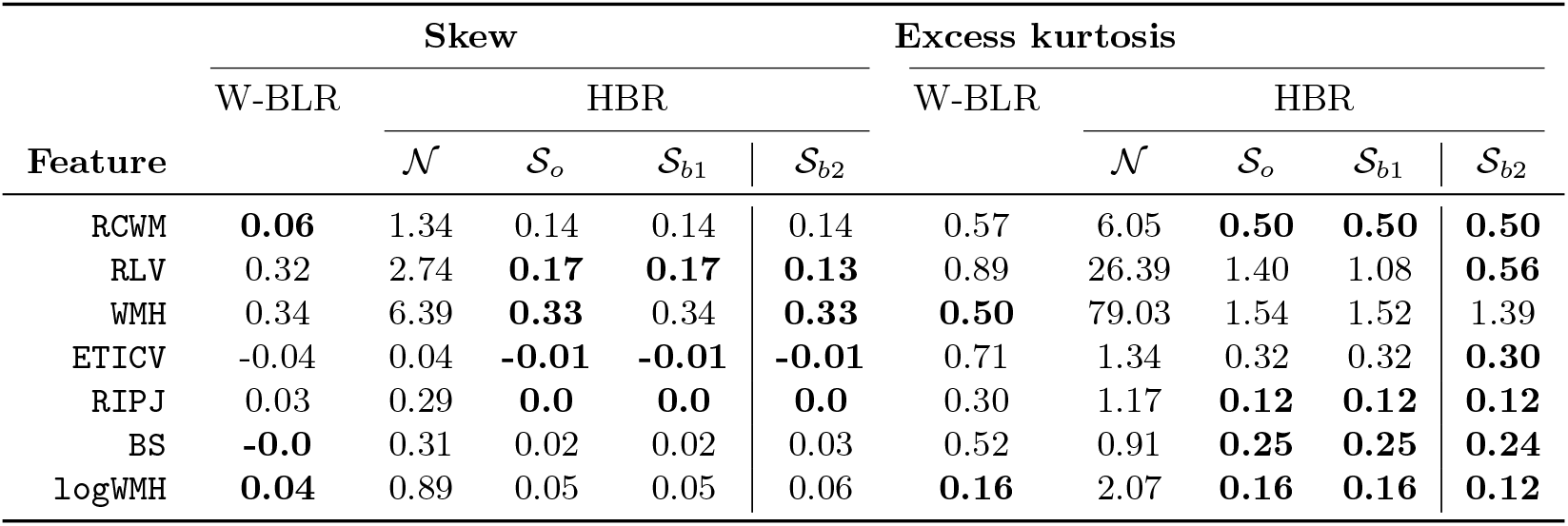
Moments of z-scores (i.e., the subject level deviations from the normative models), average of 10 folds. Smaller is better. Note that the most direct comparison is between W-BLR, 𝒮_*o*_ and *𝒮*_*b*1_. The bold figures on the left of the vertical line show the best performing method amongst these methods. The *𝒮*_*b*2_ models are more flexible by construction and have an additional random effect and phenotypes that are bold on the right side of the line are those that benefit from this additional flexibilty. ‘Hard’ phenotypes: RCWM, RLV, (log)WMH. ‘Easy’ phenotypes: ETICV, RIPJ, BS. Abbreviations: RCWM: Right-Cerebellum-White Matter; RLV: Right-Lateral-Ventricle; WMH: White-Matter-Hypointe]sities; ETICV: Estimated Total Intracranial Volume; RIPJ: interim prim-Jensen thickness (right Sulcus); BS: Brain Stem; log WMH: logWhite-Matter-Hypointensities. See also section 2.2

| Feature | Skew |  |  |  |  | Excess kurtosis |  |  |  |  |
| --- | --- | --- | --- | --- | --- | --- | --- | --- | --- | --- |
|  | W-BLR | HBR |  |  |  | W-BLR | HBR |  |  |  |
| | | $\mathcal{N}$ | $\mathcal{S}_o$ | $\mathcal{S}_{b1}$ | $\mathcal{S}_{b2}$ | | $\mathcal{N}$ | $\mathcal{S}_o$ | $\mathcal{S}_{b1}$ | $\mathcal{S}_{b2}$ |
| RCWM | **0.06** | 1.34 | 0.14 | 0.14 | 0.14 | 0.57 | 6.05 | **0.50** | **0.50** | **0.50** |
| RLV | 0.32 | 2.74 | **0.17** | **0.17** | **0.13** | 0.89 | 26.39 | 1.40 | 1.08 | **0.56** |
| WMH | 0.34 | 6.39 | **0.33** | 0.34 | **0.33** | **0.50** | 79.03 | 1.54 | 1.52 | 1.39 |
| ETICV | -0.04 | 0.04 | **-0.01** | **-0.01** | **-0.01** | 0.71 | 1.34 | 0.32 | 0.32 | **0.30** |
| RIPJ | 0.03 | 0.29 | **0.0** | **0.0** | **0.0** | 0.30 | 1.17 | **0.12** | **0.12** | **0.12** |
| BS | **-0.0** | 0.31 | 0.02 | 0.02 | 0.03 | 0.52 | 0.91 | **0.25** | **0.25** | **0.24** |
| logWMH | **0.04** | 0.89 | 0.05 | 0.05 | 0.06 | **0.16** | 2.07 | **0.16** | **0.16** | **0.12** |

The best results—results that are closest to 0—are printed in bold font but note that the most direct comparison is between W-BLR, *𝒮*_*o*_ and *𝒮*_*b*1_. The bold figures on the left of the vertical line that separates W-BLR, *𝒮*_*o*_ and *𝒮*_*b*1_ from *𝒮*_*b*2_ show the best performing method amongst these methods. The *𝒮*_*b*2_ models are more flexible by construction and have an additional random effect. Phenotypes (rows) that are bold on the right side of that vertical line are those that benefit from this additional flexibilty. For most phenotypes, *𝒮*_*o*_ or *𝒮*_*b*1_ have the best result, with a few benefiting from the additional flexibility introduced by _*b*2_, but the difference with other methods is sometimes small. The two moments need to be weighted differently as well. For instance, for the `Brain-Stem` phenotype, the W-BLR has only a marginally smaller skew than *𝒮*_*o*_, but *𝒮*_*o*_, *𝒮*_*b*1_ and *𝒮*_*b*2_ have a significantly smaller kurtosis. It is important to remember that the kurtosis is a sum of fourth powers, and the skew is a sum of third powers. Deviations thus generally have a larger effect on the kurtosis than on the skew. For the `WM-hypointensities` phenotype, the W-BLR method performs on par with the *𝒮* models regarding the skew, and better when regarding the kurtosis, indicating a better fit on those phenotypes that are also most difficult to model. The W-BLR has inherent batch effects in all aspects of the predicted percentiles, possibly explaining why it fits better to these hard phenotypes. This is further discussed in section 3.2.1. We emphasise that the comparison is not completely straightforward with the HBR models being more conservative, as described in detail in section 3.2.1. Despite that, the numbers indicate a fit that is overall better than W-BLR.

Figure 5 further illustrates the good performance of the *𝒮*_*o*_ and *𝒮*_*b*_ models relative to the *𝒩* model.

**Figure 5:**
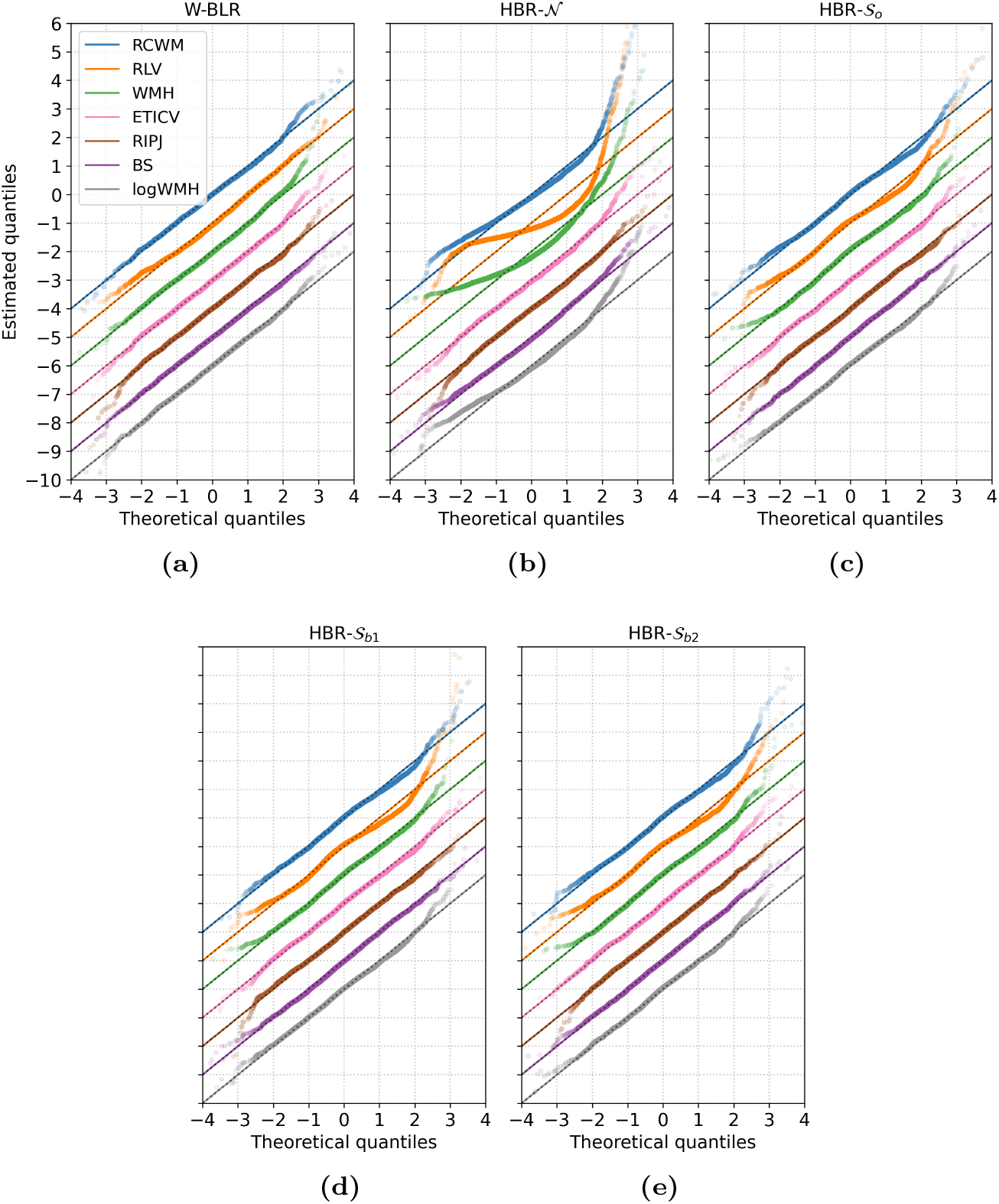
Qqplots of the estimated z-scores (i.e., the subject level deviations from the normative models) of the first test fold. Estimated z-scores of subsequent phenotypes are offset by -1 for easy comparison. For the HBR models (b), (c), (d), (e) the z-scores are computed by MAP estimation. (a) shows very straight lines, almost perfectly covering the model line. (b) shows the fit of the HBR model with Gaussian likelihood. Those phenotypes of which the residuals follow a Gaussian distribution are modeled quite well, but the limitations become very apparent when looking at the top three qqplots, which represent ‘hard’ phenotypes. The lines are very curved, and do not reflect a good fit at all. (c), (d) and (e) show a much improved fit over (b), with the plotted points laying mostly on top of the model line. Subtle differences can be found between (d) and (e), mostly in the outer centiles, where |`theoretical quantiles`| *>* 2. Abbreviations in the legend and the distinction between ‘hard’ and ‘easy’ phenotypes are specified in section 2.2.

The three hard phenotypes are not modeled well at all by the *𝒩* model. Overall the *𝒮*_*o*_ models perform comparable to W-BLR, except for `Right-Lateral-Ventricle`, where the W-BLR line looks slightly better. Note that this figure shows only a random selected cross-validation fold, whereas the statistics in Table 3 are averaged across all folds. Note also that the log transform does not by itself adequately model non-Gaussianiy in the `WM-hypointensities` phenotype, indicating that whilst it addresses non-negativity, it is still necessary to combine this transformation with a principled approach for modelling non-Gaussianity, such as afforded by the SHASH distribution.

Figure 6 visualizes the centiles of the fitted HBR models on two hard phenotypes;

**Figure 6:**
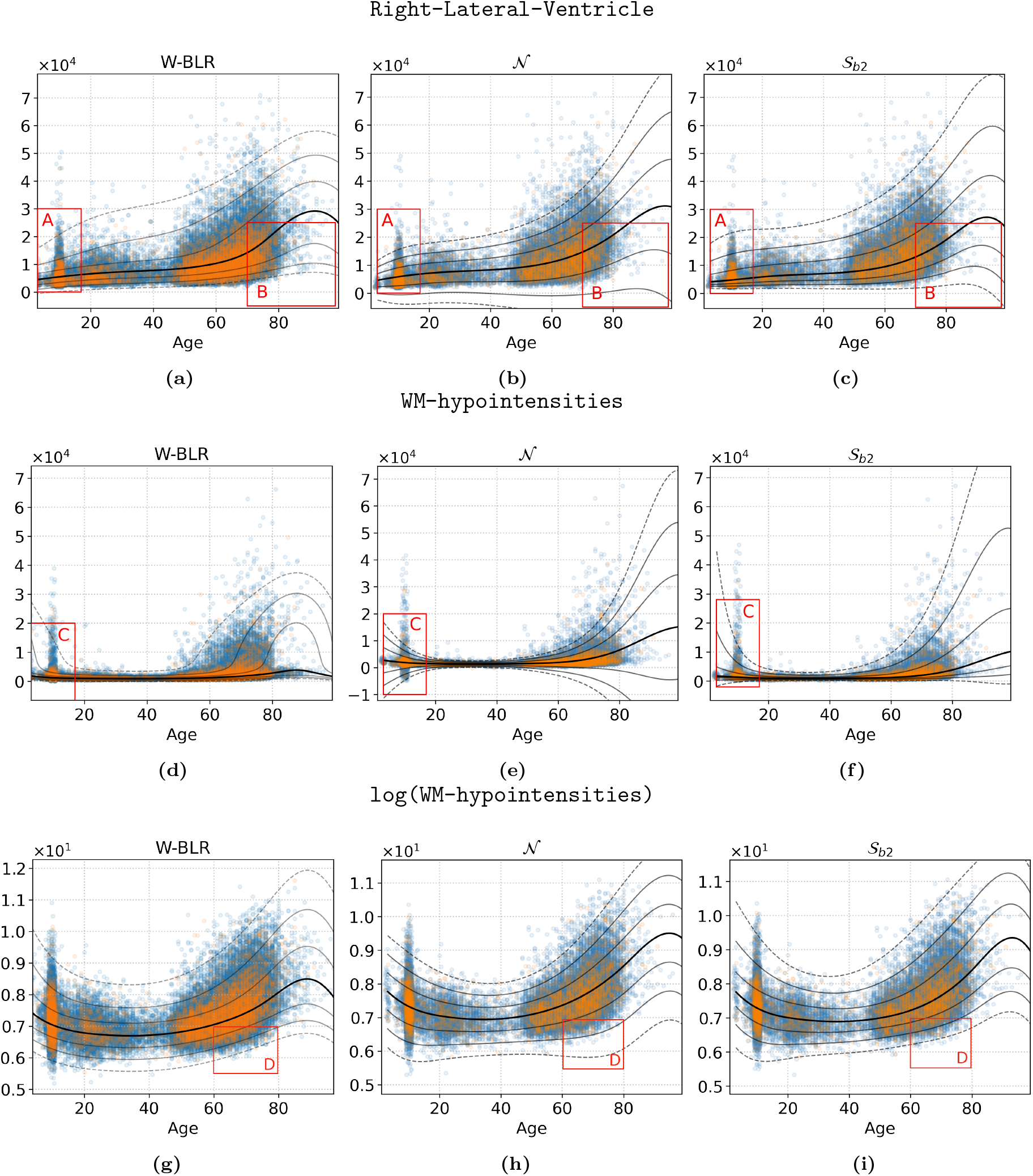
MCMC-estimated fitted centiles of `Right-Lateral-Ventricle` (top row), and `WM-hypointensities` (middle row), and log transformed `WM-hypointensities` (bottom row) by three different HBR models (columns) all ‘hard’ phenotypes). Blue markers indicate train data, the orange dots indicate the test data. The plotted data were corrected by subtracting the offset *τ*_*µ*_, learned by the corresponding model (see section Appendix B). The lines mark the median (thick line) and the first three negative and positive standard deviations in the Gaussian space, acquired by transforming the values *{*−3, …, 3*}* by equation 10. Thus, the seven plotted fitted centile lines represent the standard deviations in the Gaussian space, mapped to the SHASH space, and correspond to the 0.1’th, 2.3’th, 15.9’th, 50’th, 84.1’th, 97.7’th, and the 99.9’th percentiles. Uppercase red letters highlight three areas where the difference between the methods is apparent (see text for details).

`Right-Lateral-Ventricle`, `WM-hypointensities` and the log transformed `WM-hypointensities`. For brevity, we show only the 𝒮 _*b*2_ model, but the fitted centiles for all models and phenotypes are shown in the appendix (i.e. including 𝒮_*o*_ and 𝒮_*b*1_). Overall, this figure shows that the non-Gaussian models fit the data better. Focusing on area A, the W-BLR the 𝒮 _*b*_ model fits the data well, but the fitted centiles of the Gaussian model (b) cannot accurately model the centiles in the lower range, and many extend implausibly into the negative range (also for panel e). Focusing on area B, and realizing that this phenotype is a volume, and can therefore never be negative, the problem is obvious. Focusing on area C, we see that the W-BLR model must severely warp the input space to be able to accurately model the data at this point. For models (d) and (f), the 0.1’th fitted centile is still estimated to run into the negative domain, towards the range of the data points. This means that the data are not modeled accurately in those regions. As noted above, this motivates the application of the log transformation to the data before fitting which enforces positivity (g-i) and avoids the fitted centiles becoming negative toward the limits of range of the input variables. However, it is still necessary to model non-Gaussianity. Focussing on area D, we see that the Gaussian model applied to the log-transformed data has a poor fit in these regions, while the other two models perform well. These visualisations indicate that the SHASH transformation fits the data better and better accommodates site effects relative to a Gaussian model.

Another important aspect of the HBR method is the way it can deal with batch effects. Unlike the W-BLR method, where one-hot encoded^6^ batch vectors are appended to the design matrix (effectively treating site as a fixed effect), causing the batch effects to directly or indirectly affect the mean, variance, skew, and kurtosis, in HBR the batch effects only affect aspects of the likelihood by ecplicitly giving the parameter that controls that aspect a random effect. By having a batch-specific offset in *µ*, sampled as a deviation from a group mean, the HBR models’ predictions are ultimately indistinguishable between batches. To assess to which degree the predictions are indistinguishable between batches, we compare the Area Under the Reciever-Operator-Curve (AUC) scores [26], which vary between 0 and 1, in a 1v1 setting. Only site was used as a batch effect in our models, so the batch effects correspond exactly to the site-ids. For each combination of two sites, we compute the AUC score using the `sklearn.metrics.roc auc score` method, where we use the binary class labels as `y true`, and the z-score predictions for `y pred`. Because an AUC score of 0.5 would indicate perfect indistinguishibility, we plot the absolute deviation from the 0.5 mark in figure 7.

**Figure 7:**
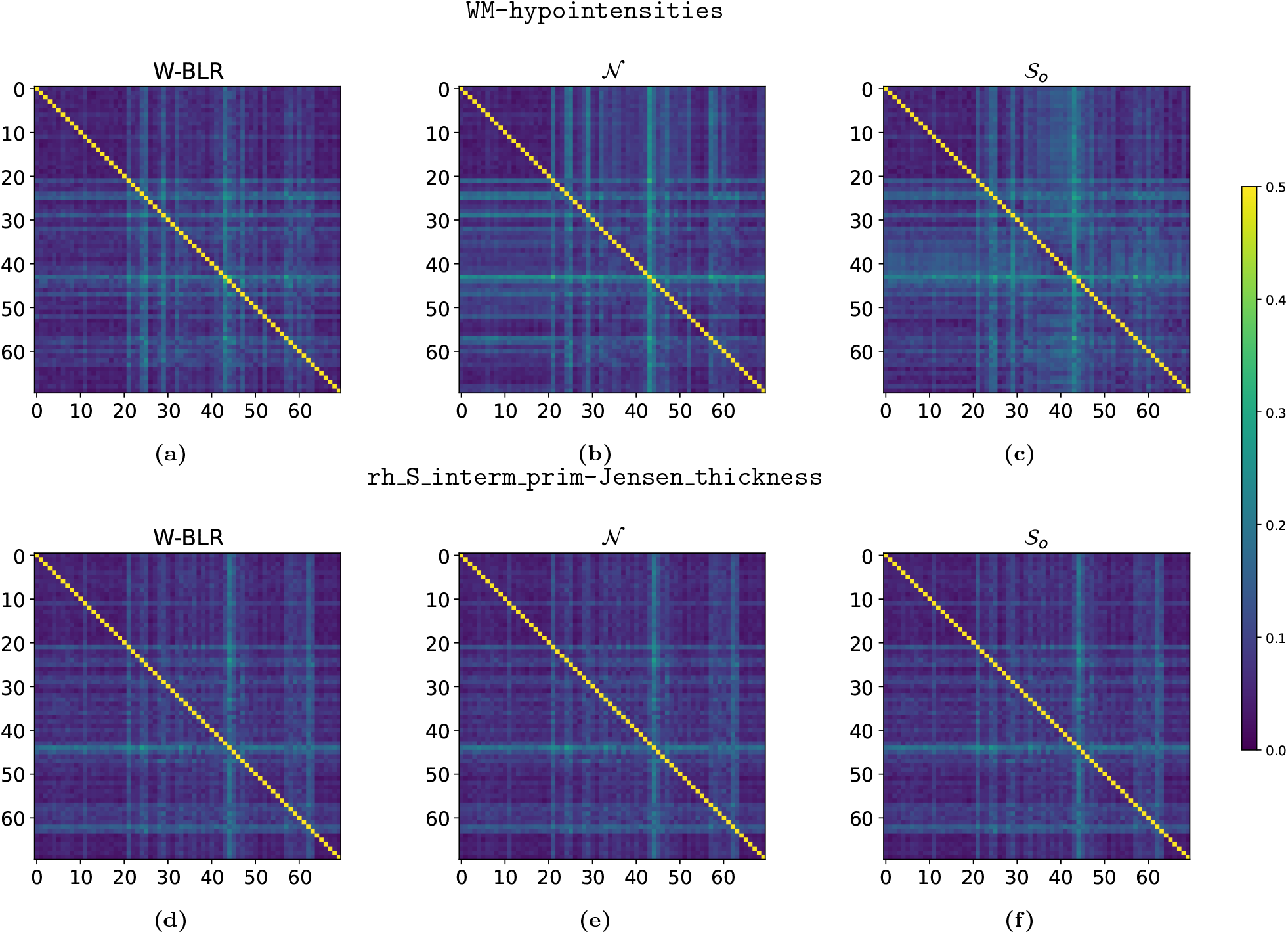
AUC scores of 1v1 site classification based on z-scores, averaged over 10 folds. The color of the x-y’th pixel represents |*a* − 0.5|, where *a* is the AUC of the binary classification problem of site x and y. Thus, the diagonal is 0.5 everywhere, which provides an upper bound to the accuracy of classification for all the for the other pixels. For off-diagonal pixels, values closer to 0 are better indicating that sites cannot be distinguished because no residual site effect remains. The top row is on z-scores of the `WM-hypointensities` phenotype, the bottom row on `rh s interm prim-Jensen thickness`. For all models, the off-diagonal terms are somewhat homogeneously close to 0. W-BLR on `WM-hypointensities` (a) seems to perform a bit better than (c) and (d), with a less distinct stripe pattern. See section 3.2.1 for a possible explanation. All other results are listed in the appendix section Appendix J, but most look similar to the plots in the bottom row; very little difference between the models is observed.

#### 3.2.1. A Subtle Difference Between HBR and W-BLR

The comparison between HBR and W-BLR seems to indicate the W-BLR should be preferred in some cases where the modeled response variable takes on an extreme shape. However, the W-BLR method and the HBR differ in the order in which they apply certain transformations, leading to a subtle difference.

The W-BLR method applies the sinh-arcsinh transformation after applying the affine transformation for mean and variance, and the HBR method applies the sinh-arcsinh transformation before applying the affine transformation for *µ* and *σ*. Because *µ* and *σ* potentially contain batch effects, and because in W-BLR, *µ* influences the shape of the distribution (as illustrated in Fig. 8), the random-effects in W-BLR indirectly influence the skew and the kurtosis. The better fit of W-BLR on the `Right-Lateral-Ventricle` and the `WM-hypointensities` data in table 3 may partly be due to this flexibility. We assume that in most cases, batch effects are not expected in the variance, skew and kurtosis as much as they are in the mean. Especially for features with a variance that is small relative to the mean, a small batch effect in the mean would not consistute a large scaling in the variance. Since the skew and kurtosis are central measures—meaning that they are computed on mean-centered data that is divided by the variance—they are not influenced by scaling or shifting transformations at all. For phenotypes derived from images from a single specific scanner may be overestimated by some constant amount, but it is less likely that the shape of the trajectory of the feature depends much on the scan location itself (although see section 4 for further discussion about this assumption). Nonetheless, a latent correlation between the site and the measurement could be present. In contrast, the HBR method provides better control because it allows modeling batch effects optionally, but not necessarily (see section Appendix B). With W-BLR, the random effects are encoded as covariates, and may therefore influence all parameters of the likelihood. The results in table 3 should be seen in light of this subtle difference. The W-BLR model may outperform HBR on some phenotypes, but the results listed under HBR are made by more conservative models, which can be controlled better in the sense that the shape parameters in HBR are implicitly regularised under the prior, whereas the shape parameters estimated by W-BLR are effectively unconstrained and can potentially more easily overfit to the data.

**Figure 8:**
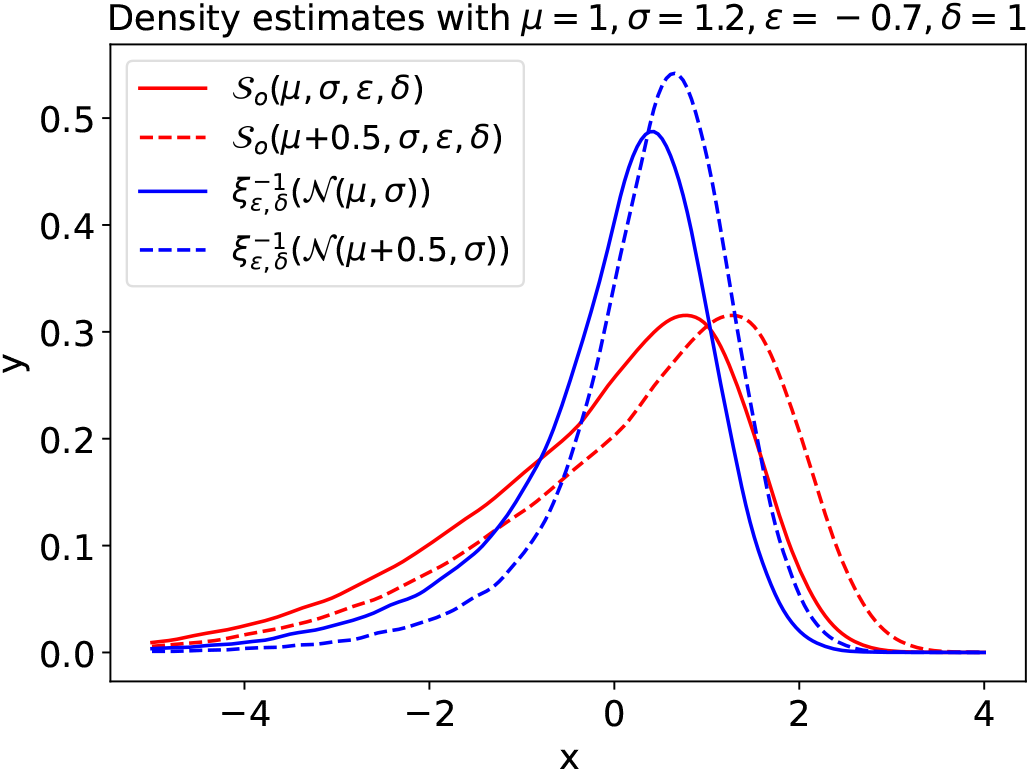
Density estimates made with 100000 samples from a standard Gaussian. For the red lines, the mean and variance (*µ* and *σ*) were added after applying the sinh-arcsinh transformation to the samples. Thus, the red lines represent a SHASH_*o*_ distribution. For the blue lines, the mean and variance were added to the Gaussian samples before applying the sinh-arcsinh transformation. The broken lines represent the result on exactly the same data, but with a slightly larger *µ*. The difference between the two red lines is limited to a shift to the right, just like one would expect when increasing *µ*. The difference between the blue lines is more than just a shift to the right, The broken blue line has a larger mode, and the general shape of the line has also changed slightly. Table 4 further illustrates this. W-BLR uses the transformation used here to create the blue line, and hence the shape of the distribution can be controlled indirectly by Φ (the design matrix, containing a basis expansion of covariates, along with the one-hot encoded batch labels in the case of W-BLR) through *µ* and *σ*, causing it to fit better to some datasets.

### 3.3. Fitting Highly Nonlinear Data With 𝒮 _b_

To demonstrate the importance of the flexibility of model 𝒮 likelihood, we take as an example a recent study in which problematic data emerged from a dimensionality reduction (see Fig. 9).

**Figure 9:**
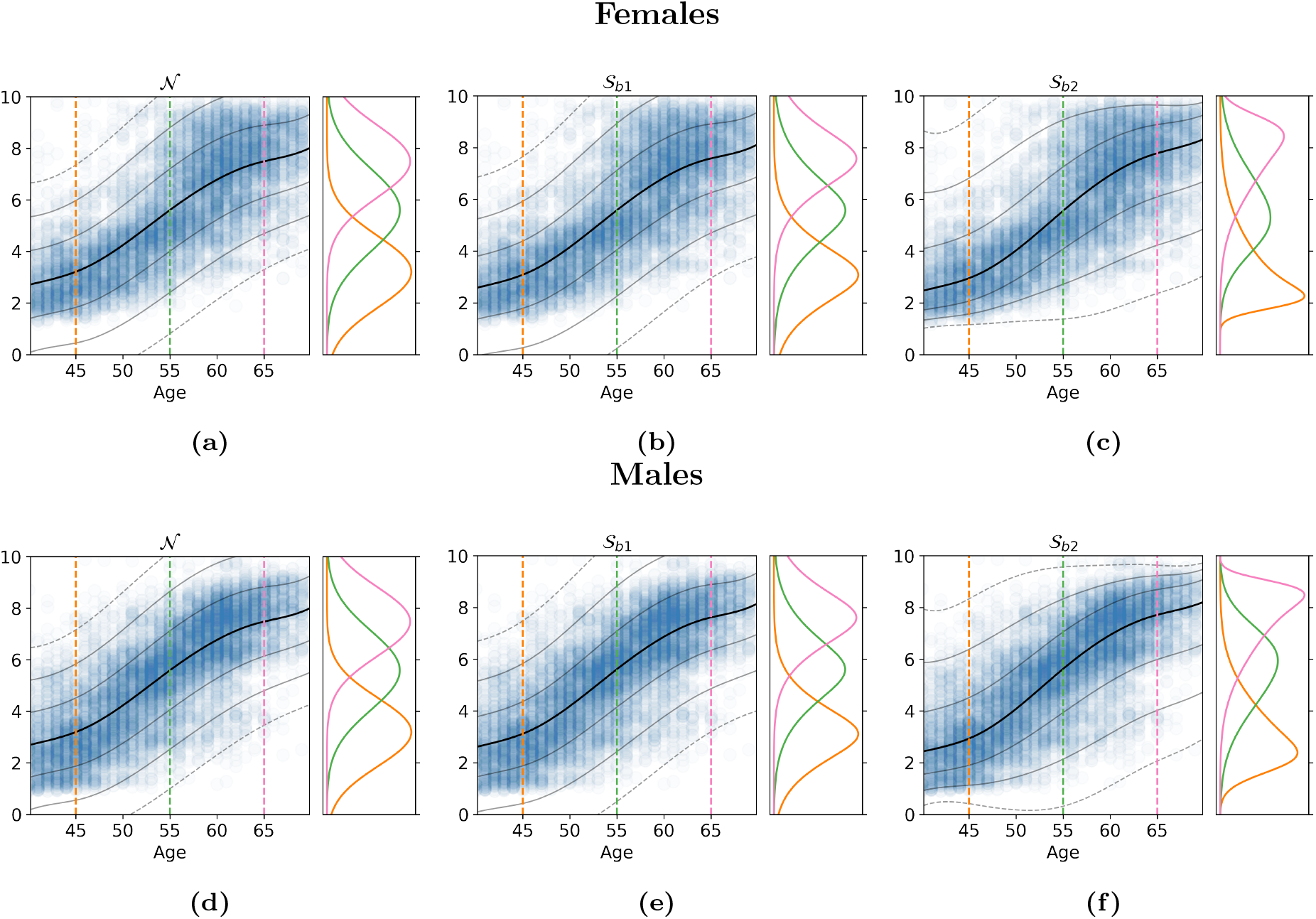
More demanding data from Zabihi et al. [12], modeled by three different HBR models. The top row is female data, the bottom row is male data. The orange, yellow, and green densities plotted on the right panels are the conditional distributions at ages 45, 55, and 65. We see that although the middle models can model for skewness and kurtosis, there is barely any difference from the Gaussian likelihoods on the left. Because the data contains roughly equal amounts of positive skew and negative skew, the best bet for a point estimate is right in the middle, at zero skew. This is reflected in the conditionals, which reflect an almost perfectly Gaussian posterior. For the models on the right, which have more flexibility in the skew and kurtosis, the conditionals change shape as they move throughout the age spectrum, truly following the data distribution.

**Table 4:** Moments of the samples used in figure 8. The first two rows differ only in the mean, by the exact amount by which *µ* was increased, whereas the bottom two rows differ entirely. This illustrates that in W-BLR, a change in *µ* changes the entire shape of the distribution.

|  | mean | variance | skew | kurtosis |
| --- | --- | --- | --- | --- |
| $\mathcal{S}_o(\mu, \sigma, \epsilon, \delta)$ | -0.24 | 2.4 | -0.96 | 0.9 |
| $\mathcal{S}_o(\mu + 0.5, \sigma, \epsilon, \delta)$ | 0.26 | 2.4 | -0.96 | 0.9 |
| $\xi_{\epsilon, \delta}^{-1}(\mathcal{N}(\mu, \sigma))$ | -0.05 | 1.27 | -1.34 | 2.76 |
| $\xi_{\epsilon, \delta}^{-1}(\mathcal{N}(\mu + 0.5, \sigma))$ | 0.37 | 0.91 | -1.31 | 3.37 |

**Table B.5:** modelling nomenclature

| Variable | Interpretation |
| --- | --- |
| $N \in \mathbb{N}$ | The number of observations, subjects, rows in the table |
| $d$ | The number of clinical covariates |
| $X \in \mathbb{R}^{N \times d}$ | Clinical covariates |
| $Y \in \mathbb{R}^{N \times 1}$ | Response variable |
| $B$ | The number of unique combinations of batch effects. So if we model 2 sexes and 7 sites, this will be $2 \times 7 = 14$ |
| $Z \in \mathbb{N}^{N \times 1}$ | A list of $N$ indices, corresponding to the combination of batch effects that apply to the data. So every $z_n \in \{1, \dots, B\}$. Not to be confused with the $z$-score. |
| $D$ | The dimensionality of the basis expansion |
| $\Phi \in \mathbb{R}^{N \times dD}$ | A design matrix derived from $X$. This can be a polynomial basis expansion, a b-spline basis expansion, or simply $X$. |

**Table F.6:** Control data, by site

| Site | # Females | # Males | Mean age |
| --- | --- | --- | --- |
| ABCD_01 | 199 | 189 | 9.89 |
| ABCD_02 | 252 | 290 | 10.07 |
| ABCD_03 | 268 | 301 | 9.88 |
| ABCD_04 | 307 | 324 | 9.81 |
| ABCD_05 | 178 | 167 | 9.89 |
| ABCD_06 | 286 | 278 | 9.94 |
| ABCD_07 | 153 | 172 | 9.87 |
| ABCD_08 | 160 | 176 | 9.95 |
| ABCD_09 | 200 | 207 | 9.96 |
| ABCD_10 | 280 | 295 | 9.86 |
| ABCD_11 | 206 | 208 | 9.81 |
| ABCD_12 | 77 | 84 | 9.88 |
| ABCD_13 | 277 | 278 | 9.81 |
| ABCD_14 | 267 | 316 | 10.19 |
| ABCD_15 | 178 | 217 | 9.90 |
| ABCD_16 | 414 | 507 | 9.90 |
| ABCD_17 | 267 | 290 | 9.81 |
| ABCD_18 | 160 | 181 | 9.90 |
| ABCD_19 | 272 | 262 | 10.06 |
| ABCD_20 | 320 | 320 | 10.05 |
| ABCD_21 | 222 | 269 | 9.91 |
| ABIDE_GU | 27 | 27 | 10.42 |
| ABIDE_KKI | 65 | 122 | 10.29 |
| ABIDE_NYU | 28 | 107 | 14.40 |
| ABIDE_USM | 3 | 56 | 22.07 |
| ADD200_KKI | 27 | 34 | 10.25 |
| ADD200_NYU | 20 | 18 | 10.21 |
| AOMIC_1000 | 483 | 445 | 22.84 |
| AOMIC_PIP02 | 120 | 89 | 22.18 |
| ATV | 17 | 60 | 22.67 |
| CIN | 22 | 44 | 47.89 |
| CMI_CBIC | 73 | 126 | 11.19 |
| CMI_RU | 125 | 251 | 10.53 |
| CMI_SI | 38 | 68 | 11.01 |
| CNP-35343.0 | 43 | 47 | 31.94 |
| CNP-35426.0 | 11 | 9 | 29.50 |
| COI | 78 | 46 | 51.86 |
| HCP_A_MGH | 86 | 85 | 59.75 |
| HCP_A_UCLA | 71 | 53 | 53.32 |
| HCP_A_UM | 120 | 84 | 61.59 |
| HCP_A_WU | 112 | 66 | 58.83 |
| HCP_D_MGH | 109 | 107 | 13.78 |
| HCP_D_UCLA | 62 | 65 | 14.13 |
| HCP_D_UM | 85 | 71 | 13.26 |
| HCP_D_WU | 75 | 79 | 13.97 |
| HCP_EP_BWH | 2 | 6 | 23.09 |
| HCP_EP_IU | 11 | 14 | 23.90 |
| HCP_EP_MGH | 3 | 8 | 27.61 |
| HCP_EP_McL | 4 | 9 | 25.15 |
| HKH | 17 | 12 | 45.41 |
| HRC | 36 | 13 | 41.69 |
| HUH | 38 | 29 | 34.74 |
| KCL | 25 | 16 | 34.12 |
| KTT | 41 | 87 | 31.03 |
| KUT | 66 | 93 | 36.50 |
| Oasis2 | 127 | 58 | 76.88 |
| Oasis3 | 631 | 921 | 69.78 |
| SWA | 15 | 85 | 28.48 |
| SWU_SLIM_ses1 | 318 | 230 | 20.07 |
| UCDavis | 62 | 74 | 3.12 |
| UTO | 106 | 96 | 35.19 |
| cam | 329 | 318 | 54.19 |
| delta | 18 | 31 | 50.44 |
| ds001734 | 60 | 48 | 25.54 |
| ds002236 | 38 | 48 | 11.49 |
| ds002330 | 37 | 29 | 26.62 |
| ds002345 | 131 | 76 | 21.69 |
| ds002731 | 28 | 31 | 21.25 |
| ds002837 | 42 | 44 | 26.73 |
| hcp_ya | 606 | 507 | 28.80 |
| ixi | 313 | 245 | 48.71 |
| nki | 308 | 174 | 42.63 |
| pnc | 701 | 677 | 14.21 |
| top | 134 | 158 | 34.57 |
| ukb-11025.0 | 12944 | 12042 | 62.98 |
| ukb-11027.0 | 5438 | 4534 | 64.39 |

**Table F.7:**
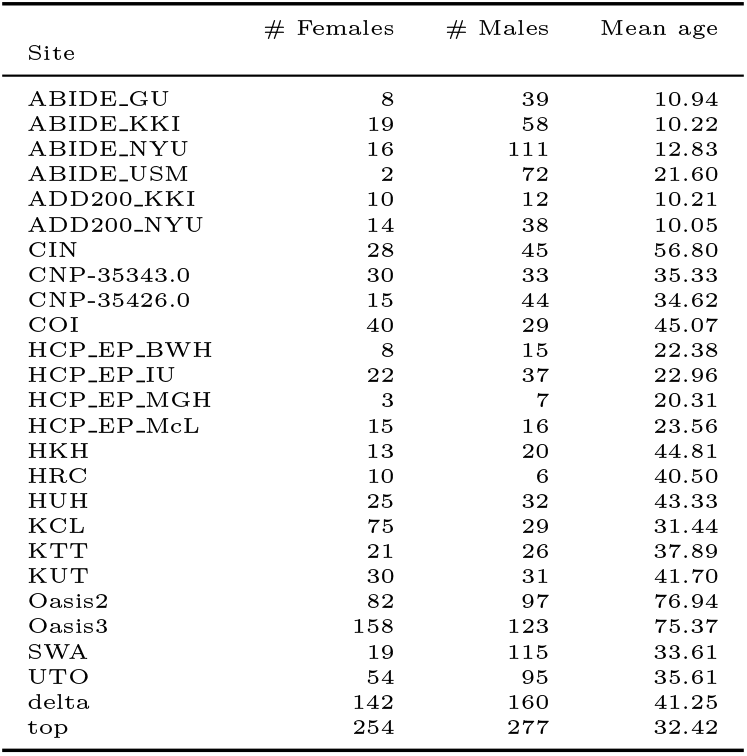
Patient data, by site

**Table F.8:** Patient data, by group

| Group | # Females | # Males | Mean age |
| --- | --- | --- | --- |
| AD | 142 | 94 | 75.75 |
| ADHD | 40 | 71 | 17.60 |
| ASD | 61 | 389 | 19.08 |
| BD | 130 | 106 | 34.67 |
| Converted | 24 | 13 | 79.75 |
| Demented | 58 | 84 | 76.21 |
| MCI | 16 | 29 | 73.40 |
| MDD | 260 | 239 | 40.26 |
| MDD_motar | 30 | 21 | 35.50 |
| Other | 15 | 26 | 34.21 |
| Pain | 25 | 39 | 55.28 |
| Patient | 72 | 98 | 30.01 |
| Patient_motar | 32 | 38 | 40.20 |
| SZ | 131 | 225 | 34.85 |
| Stroke | 3 | 6 | 67.66 |
| otherpsychosis | 32 | 52 | 29.76 |
| schizoaffective | 23 | 10 | 35.69 |
| schizophreniform | 11 | 12 | 25.43 |

In [12], fMRI data was compressed using a autoencoder [22], and then further compressed using UMAP dimensionality reduction [23]. The result was a 2d latent representation of brain activation patterns, which followed a somewhat problematic, highly nonlinear, distribution. Postively skewed on the lower end, and negatively skewed on the higher end of the age range, this representation can not be modeled by HBR with a Gaussian likelihood, and W-BLR is also not designed to deal with these kinds of effects. If we take 𝒮_*b*1_, we find that we run into the limits of that model as well, because it assumes a fixed skew and kurtosis throughout the age range. With model 𝒮_*b*2_ we have the flexibility to model heterogeneous skew and kurtosis, and Fig. 9 shows that this flexibility is absolutely essential for modelling this phenotype.

### 3.4. Computational Complexity

We provide an indication here about the computational complexity for the time required to estimate the different variants of our models. Specifically, for the lifespan normative models presented in section 2.2 above the mean [stdev] time averaged over 10 folds and specified as hours:minutes). This measures the time required to draw 1000 samples after 500 samples burn in was: 𝒩 = 2:19 [0:36], 𝒮_*o*_ = 6:29 [1:00], 𝒮_*b*1_ = 6:54 [0:31] and 𝒮_*b*2_ = 27:26 [1:27]. Another important question in the use of MCMC samplers is determining the number of samples to acquire (i.e. the chain length). We provide a simple evaluation to determine whether 1000 samples after burn-in is sufficient. To achieve this we compare the empirical mean and variance of different parameters acros chains having 500, 1000 and 1500 samples, post burn in. We also show Monte Carlo estimates for a representative set of parameters in Figure J.36. This shows that the parameters are effectively identical, with perhaps a slight reduction in the Monte Carlo estimates of the variance for some parameters for the short chains. Overall, this provides confidence that the chain length we employ is sufficient for reliable inferences.

## Discussion

In this work, we proposed a flexible method for modelling non-Gaussianity in normative models based on the SHASH distribution. We proposed a novel reparameterisation and developed an efficient MCMC sampling approach for these models. We first applied this method to a large neuroimaging dataset and show competitive performance with respect to competing methods, whilst providing full Bayesian inference and full distributions over all model parameters. Then, we applied this method to a highly challenging problem where we aim to model a set of learned parameters from a highly non-linear model (derived from an autoencoder). In this case the classical methods are not able to satisfactorily model the distribution of the brain phenotype, while the proposed method can.

Normative modelling aims to find centiles of healthy variation in neuroimaging phenotypes, much like growth charts in pediatric medicine. Many of those phenotypes follow complex nonlinear, asymmetric distributions, so for a normative model to be generally applicable, it has to be able to model features like skewness and kurtosis. Our extension builds upon the HBR framework for Normative modelling introduced by Kia et al. [5], which retrieves a distribution over model parameters given a prior distribution over the parameters and a likelihood over the data. The original paper reported only on results where a Gaussian likelihood was used, but the framework is flexible enough to support any parameterized likelihood. Here we adapted the HBR framework to make use of a different one, namely the SHASH likelihood, which has additional parameters roughly modelling for skew and kurtosis.

The basic SHASH distribution in equation 3 has no parameters for location and scale, but those can be added, resulting in the SHASH_*o*_ (*𝒮*_*o*_) distribution, as in equation 5. We also introduce a reparameterization called the SHASH_*b*_ (*𝒮*_*b*_) distribution, which aims to remove the severe correlations by applying an affine tranformation for location and scale to a standardized SHASH distribution.

Our results on a large neuroimaging dataset show that the *𝒮*_*b*_ reparameterization in some cases converges better and more reliably than the *𝒮*_*o*_ distribution. For example, in the analysis of the highly challenging data, shown in Fig. 9, the *𝒮*_*o*_ model did not converge. However, for most phenotypes the convergence is similar to the original variant. A more important benefit of this reparameterisation is that the parameters of the distribution are more interpretable, as shown schematically in Fig. 1 and empirically in Fig. 2. We note that the *𝒮*_*b*_ distribution is isomorphic to *𝒮*_*o*_—the two can model exactly the same set of distributions— which might not give a preference for one distribution over the other in the context of HBR. We also observed a similar sampling performance (at least in terms of convergence) for both. However, the *𝒮*_*o*_ parameterisation could in some circumstances be preferred over *𝒮*_*b*_, (e.g. for ‘easy’ phenotypes with near-Gaussian shapes) if parameter interpretation is not of primary importance, although the latter achieved slightly better performance in our experiments. However, as noted above, this *𝒮*_*o*_ model did not converge for the most difficult nonlinear phenotype we considered, in which case *𝒮*_*b*_ is preferred.

We showed that HBR with a *𝒮*_*b*_ likelihood performs equivalently or in some cases slightly better than Warped Bayesian Linear Regression (W-BLR) [8] on most datasets, in terms of Gaussianity of the predicted deviation scores, although W-BLR predictions are better on specific pathological phenotypes. We argue that the W-BLR method fits better to that data due to the fact that batch effects are effectively encoded as a fixed effect and can therefore potentially affect all parameters of the likelihood, which is not always desired (See Fig. 8). The W-BLR method has no option to disable this flexibility, as it is inherent to the method. This is why we argue that the HBR method should be preferred as it supports enabling or disabling random effects in each parameter of the likelihood as a modelling choice. Moreover, the W-BLR method is unable to model the most complex phenotype we evaluated (i.e. derived from the latent representation of an autoencoder) because it does not provide the ability to sufficiently control the skewness and kurtosis across the range of the covariates. In other words, HBR provides more flexible parametric control over the shape of the distributions that the approach is able to model.

We have shown that HBR with *𝒮*_*b*_ likelihood is able to model highly non-Gaussian distributions with intricate trajectories in several parameters in a way that is impossible to replicate with existing techniques. This indicates that HBR will possibly play an important role in pushing the field of Normative modelling forward.

The computational complexity of the likelihood may be the most important limitation, especially for the *𝒮*_*b*2_ variant. It must be noted that the computation time is significantly reduced if fewer batch effects are present. We hope that future releases of software packages like `pymc` will help reduce the required computational time.

Lastly, all our work is freely available to be used and extended, in the open-source `pcntoolkit` package.

### 4.1. Future Work

#### 4.1.1. Transfer Learning

In the current work, we have not applied the method in a transfer learning setting, but the methods for doing so are described in [5] and the updates we provide to the `pcntoolkit` package support this. For transfering a learned model to a new site, one would have to create a factorized approximation of the posterior by using the MCMC samples retrieved earlier. Restricting ourselves to a known distributional form, this can easily be done by standard optimization techniques on the log likelihood. The factorized posterior is then fixed, and can be used as an informed prior for MCMC sampling with data from a new site. The samples retrieved in this manner can then be used for approximation of z-scores or any other downstream analysis.

#### 4.1.2. Variational Inference

We have shown the value of the *𝒮*_*o*_ and *𝒮*_*b*_ likelihoods in the context of MCMC sampling. The reparameterization removes much of the correlation in the posterior that is problematic for sampling methods, leading to more stable results. Nonetheless, the MCMC sampling procedure is still very computationally demanding, especially on large datasets with many batch effects. A variational approach [28] may now be a viable solution to this problem. In the HBR context, the *𝒮*_*b*_ reparameterization may be particularly beneficial in this regard because it removes dependencies between the parameters, such that the posterior can be approximated by a factorized distribution. Variational inference is a fast approximation, and MCMC is a slow but (asymptotically) exact method. We believe that in some cases a fast approximation suffices, so variational approximation could be a suitable extension to the present work.

#### 4.1.3. Generalisations to more expressive noise distributions

The extension presented here is just the tip of the iceberg of Hierarchical Bayesian modelling. This extension was developed for modelling skewness and kurtosis, and while we showed that this can fit a wide range of distributions, the flexibility of the SHASH distribution is still limited. Specifically, the limitations in tail thickness are easily read from plot 1d. Mixture models could further help increase the expressive power of the HBR method. One could design a mixture of a single SHASH_*b*_ (or simply a Gaussian, which might be easier to fit) and two Gamma distributions, one left-tailed and one right-tailed, conveniently located as to act as surrogate tails. The central *𝒮* component would be dedicated to learning the shape of the bulk of the data, while the Gamma distributions absorb the largest outliers. With the current version of the `pcntoolkit`, this extension is now within reach, but we leave this for future work.

#### 4.1.4. Batch Effects

Multidimensional batch effects are currently implemented as a *D* dimensional matrix in the `pcntoolkit`, containing one value per combination of batch effects. The number of values grows exponentially with the number of batch effects. An alternative approach is to model a separate vector for each batch effect, and to learn parameters linking the vectors together. So instead of having *θ*_*i,j*_ ∼ *𝒩* (*µ*_*i,j*_, *σ*), one could have 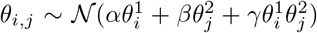. The number of required linking parameters grows as *n*(*n* + 1)*/*2 ∈𝒪 (*n*^2^), much better than the current 𝒪 (*d*^*n*^), where *d* is the number of unique values for a batch effect. We expect that the correlation between batch effects is not sufficient to justify the approach, and we believe the most important effects in the batches can be modeled this way. In addition, throughout this work we have assumed that the batch effects are principally evident in the mean, and observed empirically that this provided reasonable performance. It is conceivable, however, that batch effects could also be evident in the variance. Whilst the original Gaussian HBR method can easily handle batch effects in the variance, the use of the SHASH distribution complicates this somewhat; while we could add batch effects to the variance as well, this might interact with the shape and/or heteroskedasticity of the resulting model. We therefore leave this for future work.

## 5. Conclusion

We have extended a method for federated normative modelling of neuroimaging data to support non-Gaussian attributes such as skew and kurtosis. To this end, we have introduced a 4-parameter reparameterization of the SHASH distribution, along with an accompanying software implementation. We have demonstrated that our method performs equivalently or in some cases slightly better than a baseline method on simple and complex data, thereby showing that this method can push the field of normative modelling forward.

## Appendix A. Sinh-Arcsinh transformation

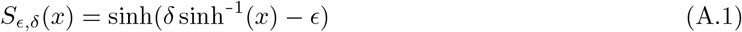

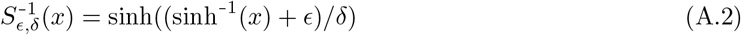

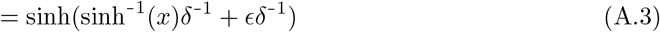

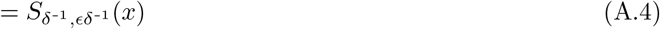

## Appendix B. Model Specification

Every phenotype has its own defining characteristics. Some may be severely skewed, while others appear to be nicely Gaussian. We may see strong correlations between age and variance (heteroskedasticity) in one, while another seems to retain a consistent variance throughout the entire age spectrum. Batch effects like sex-specific differences, or site-specific noise characteristics may or may not be present in the data. Characteristics like these can be learned by a sufficiently flexible model. However, Occam dictates that we should prefer a simple model over a complex model, if they perform equally well. A model is therefore optimally defined if it has enough flexibility to capture the parameters of interest, whilst being constrained in every other way. An ideal modelling framework would allow specifying constraints on all aspects of the model, be flexible enough to support a wide range of models, while still remaining concise. Here a model definition framework is proposed that allows us to concisely write down a wide range of different models. This framework will simplify the later sections of this report. In addition, the framework introduced here aligns with the implementation of the source code in the `pcntoolkit` software package. The graphical notation used here is slightly adapted from [16] chapter 8, where here we indicate the range and name of the iterator variables more explicitly in the plates, i.e. *N* becomes *n* ∈ { 1, …, *N*} . For the remainder of this section we will uphold the nomenclature in Tab. B.5. Without loss of generality, let us now focus on modelling a single parameter, *θ*. For more complex distributions that take more parameters, like almost any distribution, we just apply what follows to every parameter separately.

### Appendix B.1. Constant θ

Here we discuss simple models where *θ* is constant for the whole spectrum of *X*, save batch effects.

#### Model ℳ_1_

Let *ℳ*_1_ represent the case where *θ* does not depend on *X* or *Z*, i.e. it takes a single value for all *y*_*n*_. The only dependency of *θ* in ℳ_1_ is on the (set of) hyperparameters *α* through its prior distribution *p*:

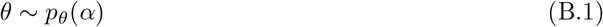

and Fig. B.10 shows the graphical model.

**Figure B.10:**
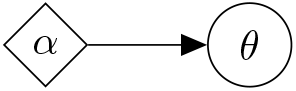
Graphical model for ℳ_1_

**Figure B.11:**
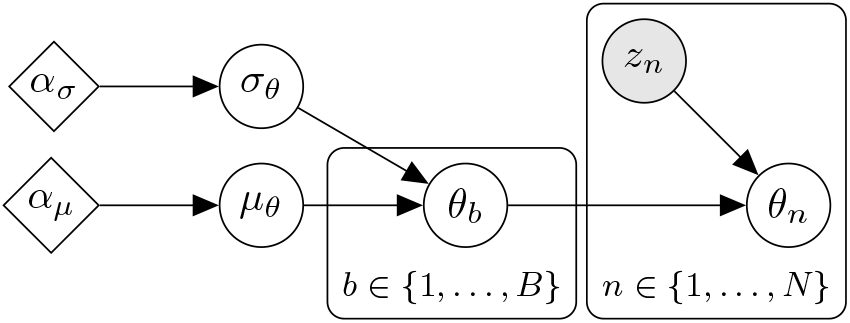
Graphical model for ℳ_2*a*_

**Figure B.12:**
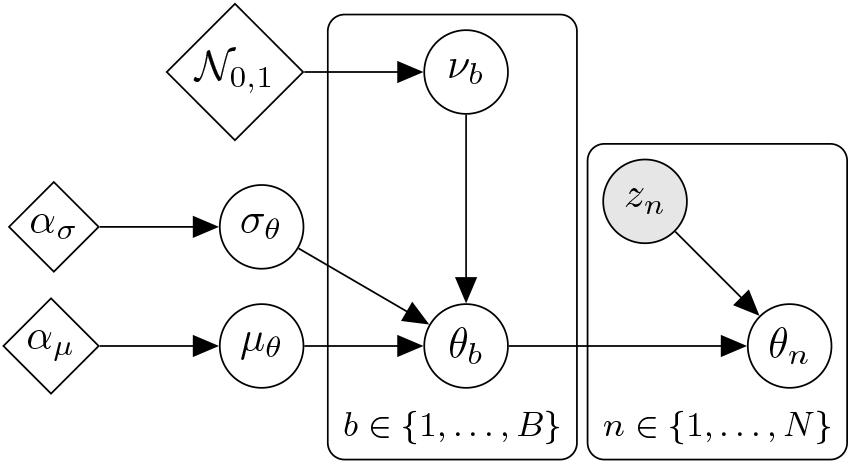
Graphical model for ℳ_2b_

**Figure B.13:**
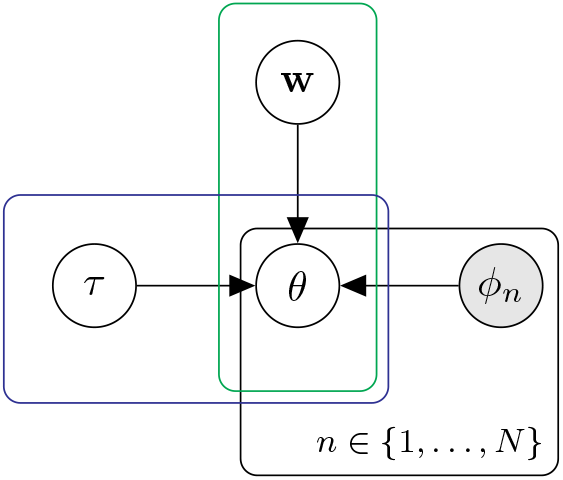
Graphical model for ℳ_3.∗.∗_. The green and blue box are added to emphasize that **w** and *τ* may be modeled with batch effects, resulting in multiple values for *θ*.

**Figure G.14:**
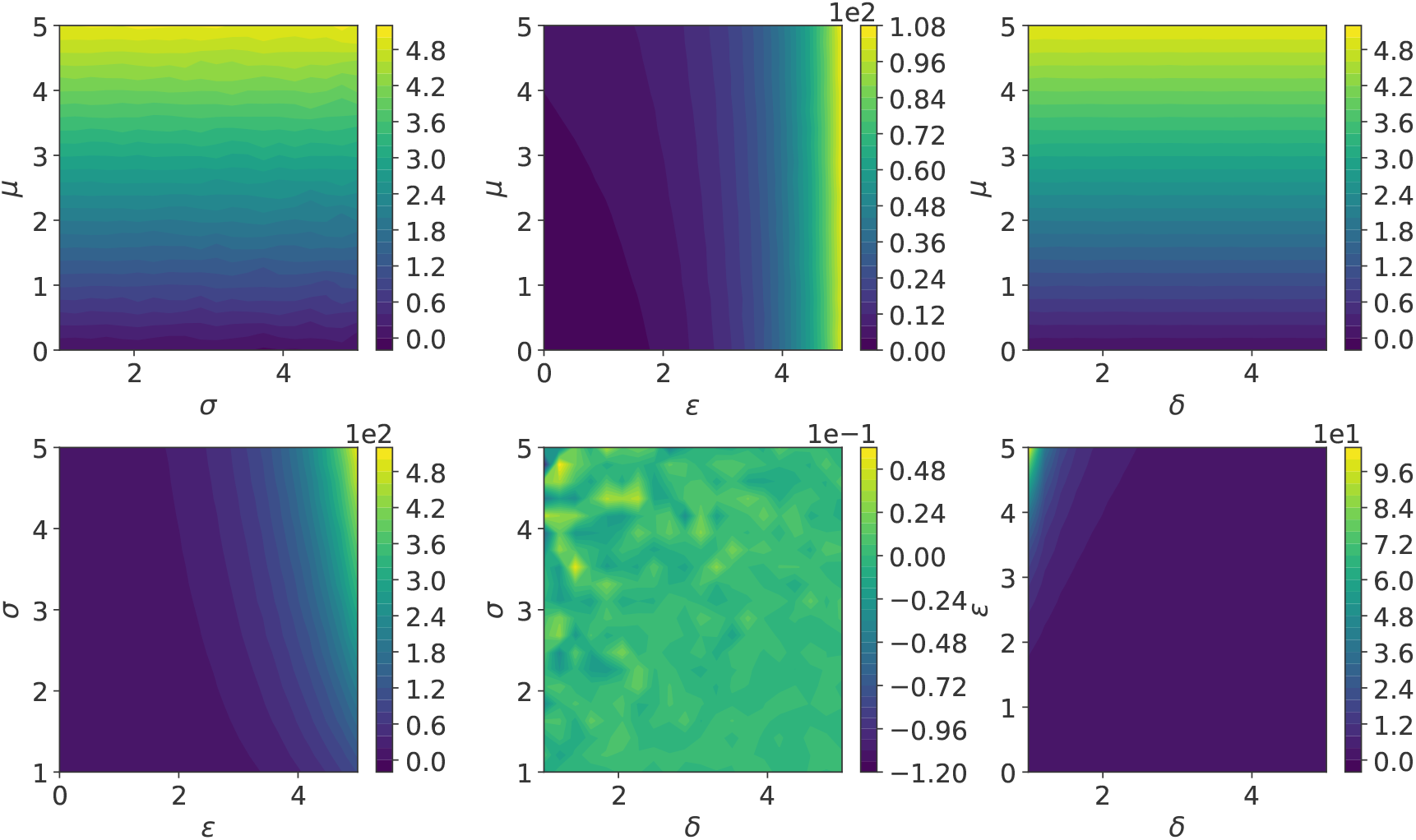
SHASH_o_ first moment *E*[*x*]

**Figure G.15:**
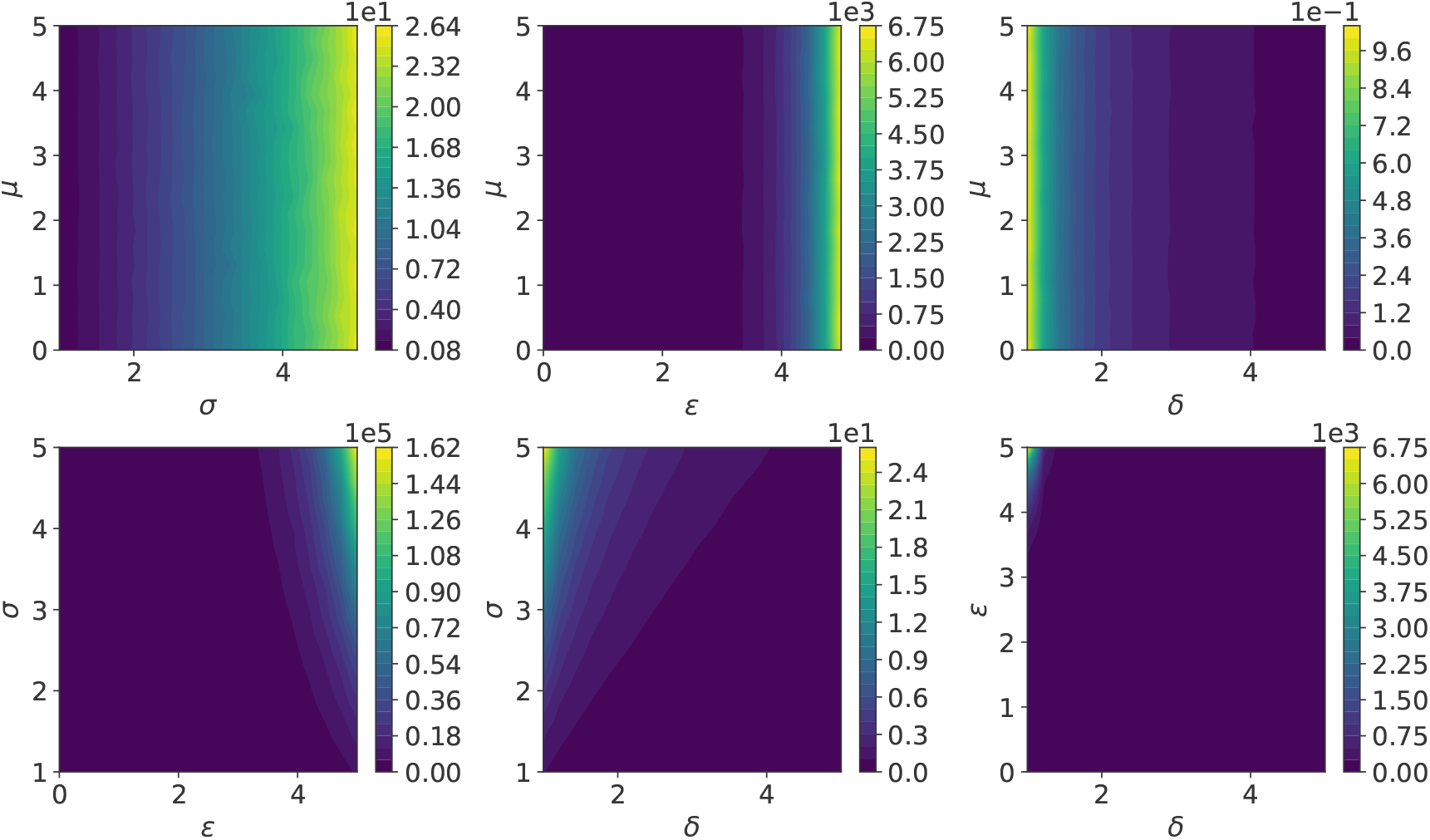
SHASH_*o*_ second moment *E*[(*x* − *µ*)^2^]

**Figure G.16:**
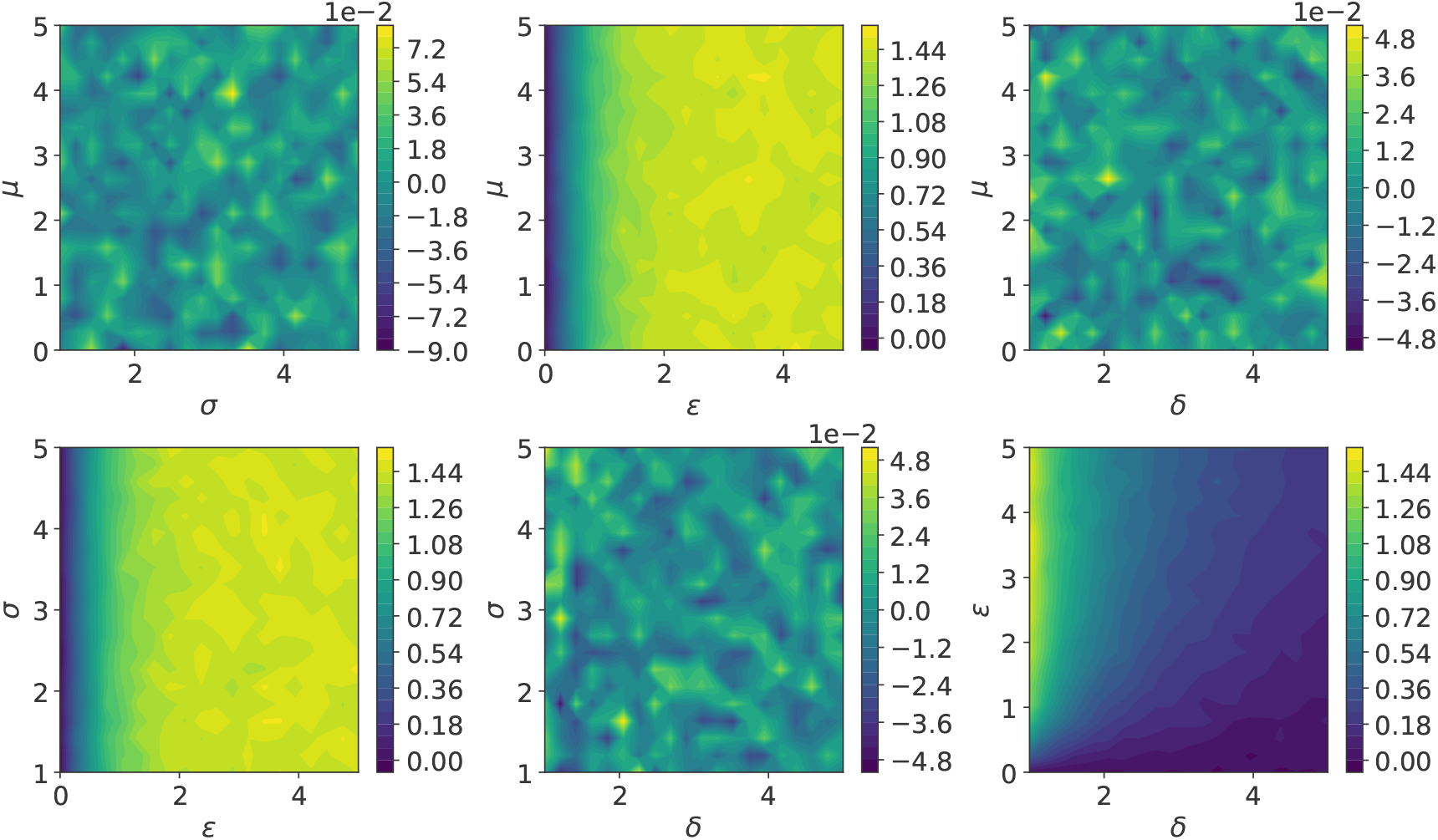
SHASH_o_ third moment *E* [((*x* − μ)/ σ)^3^]

**Figure G.17:**
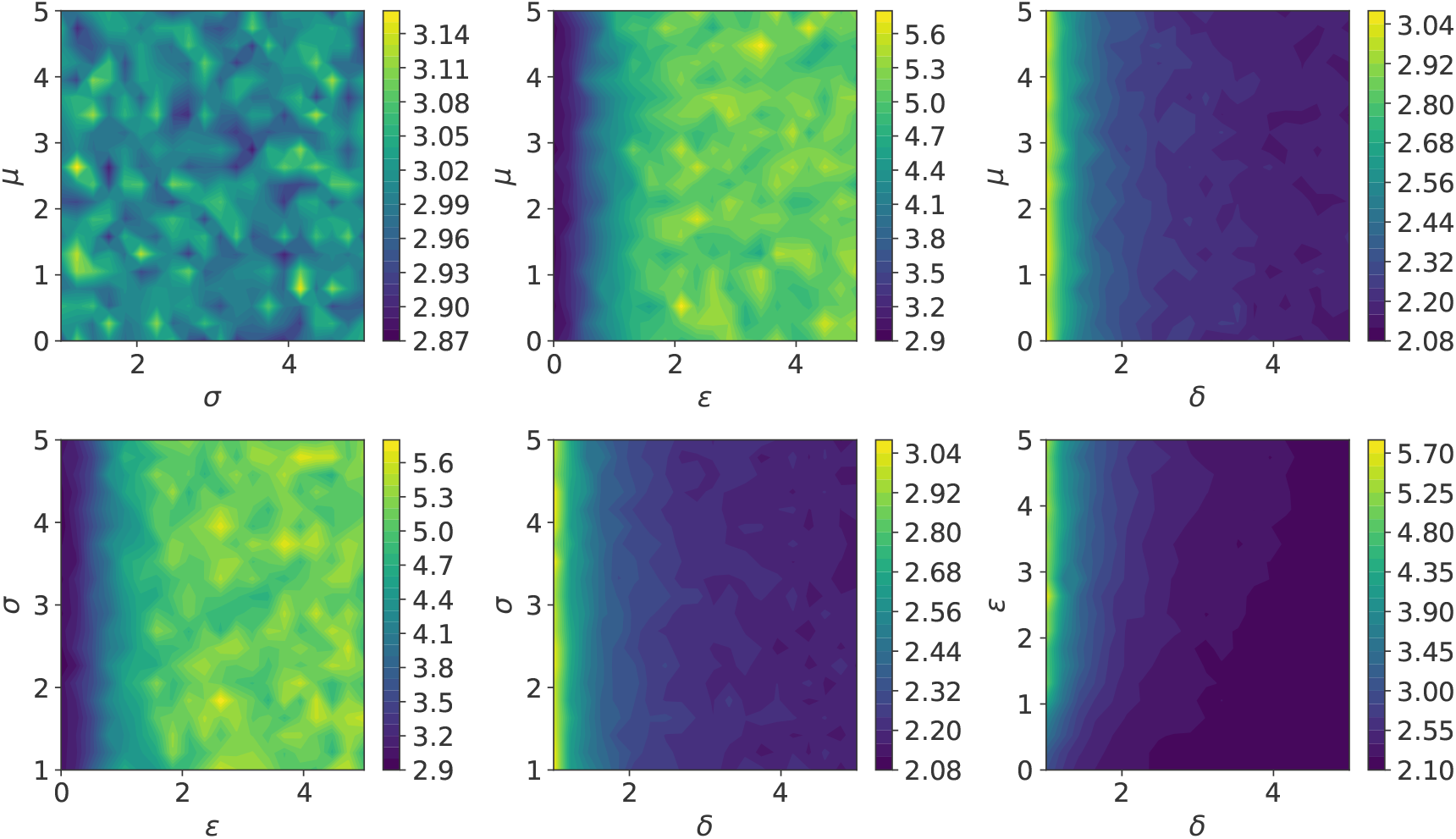
SHASH_*o*_ fourth moment *E*[((*x* − *µ*)*/σ*)^4^]

**Figure H.18:**
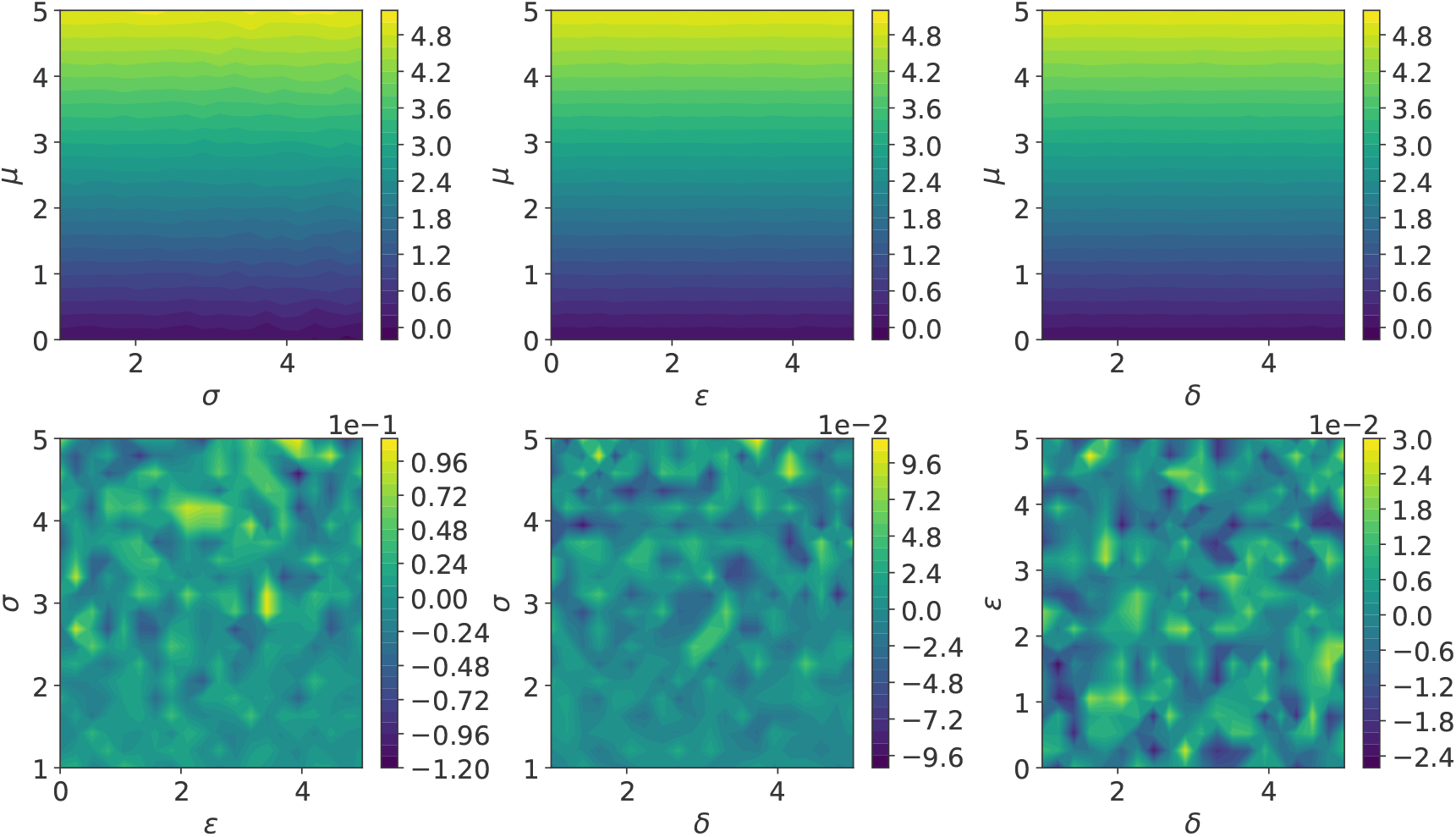
SHASHb first moment *E*[*x*]

**Figure H.19:**
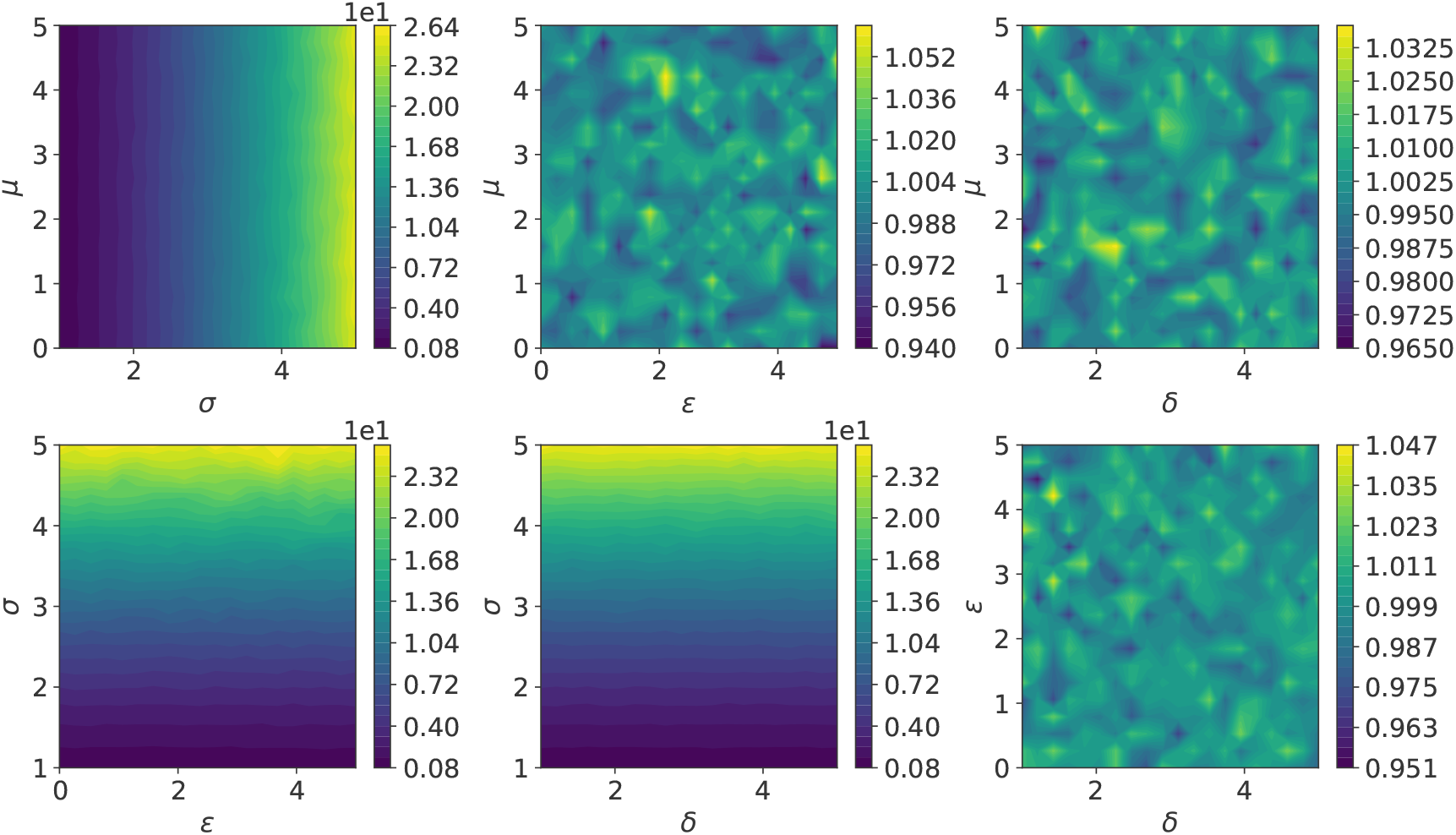
SHASH_*b*_ second moment *E*[(*x* − *µ*)^2^]

**Figure H.20:**
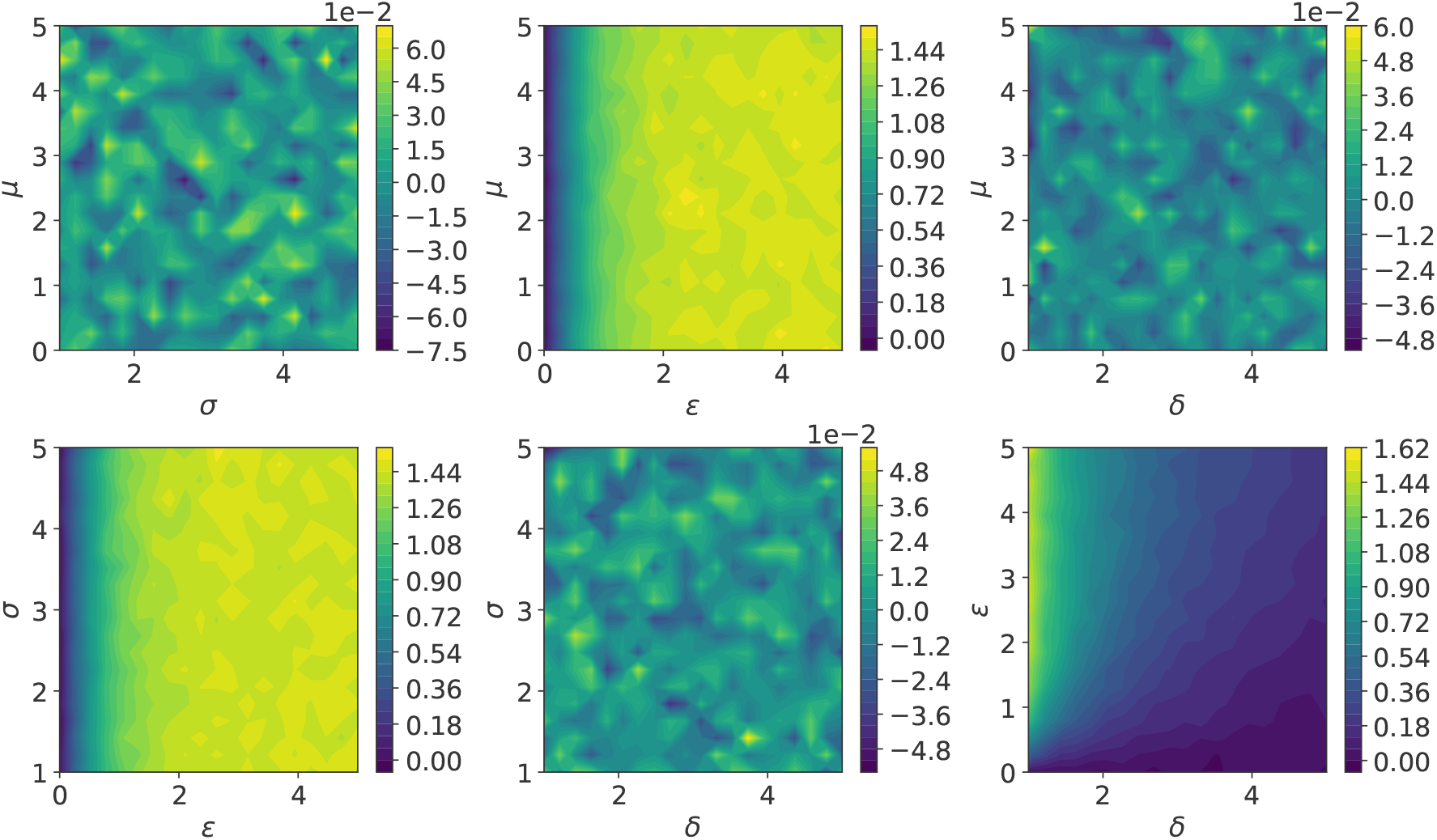
SHASHb third moment *E*[((*x* − μ)/σ)^3^]

**Figure H.21:**
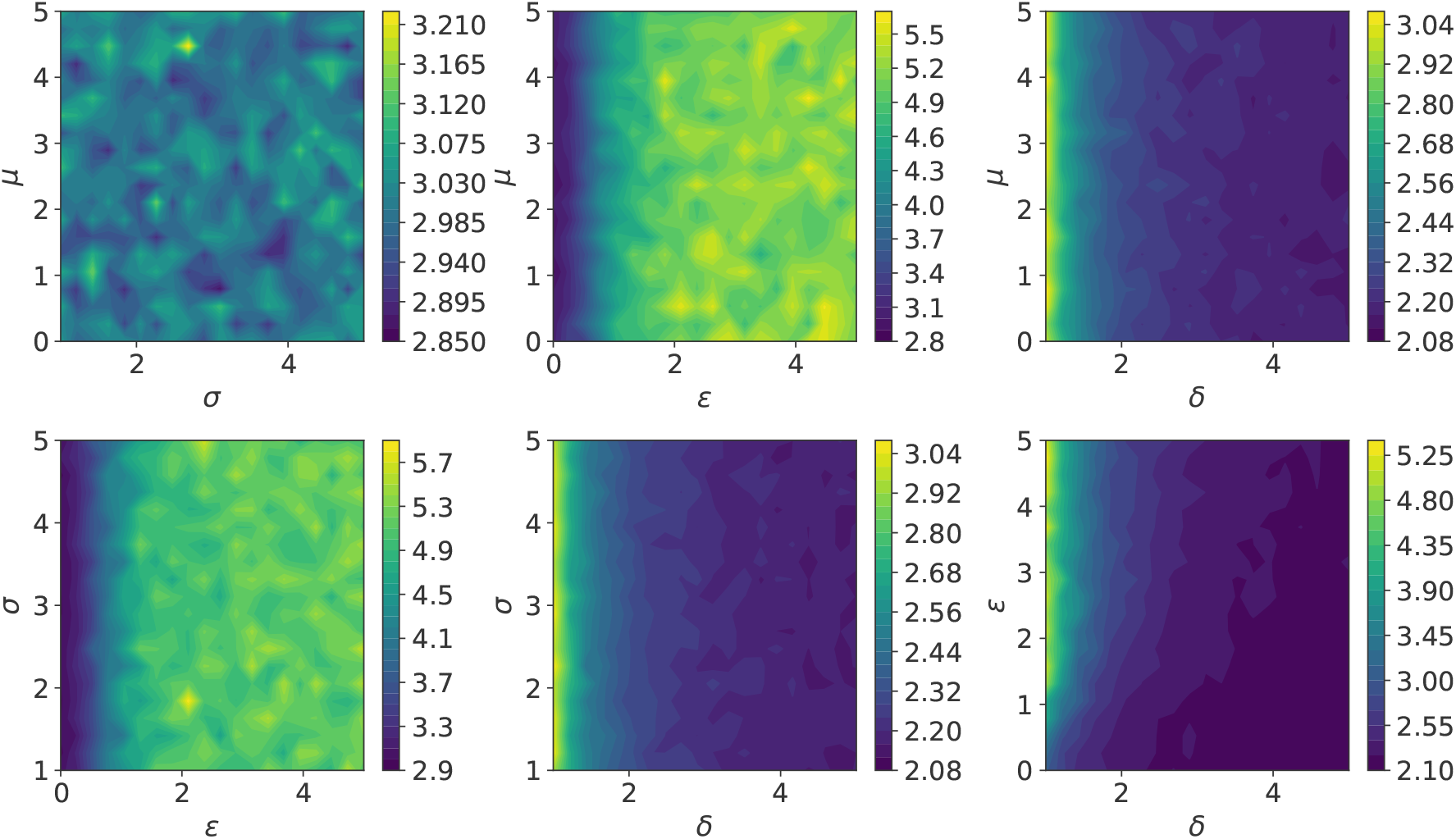
SHASH_*b*_ fourth moment *E*[((*x* − *µ*)*/σ*)^4^]

**Figure I.22:**
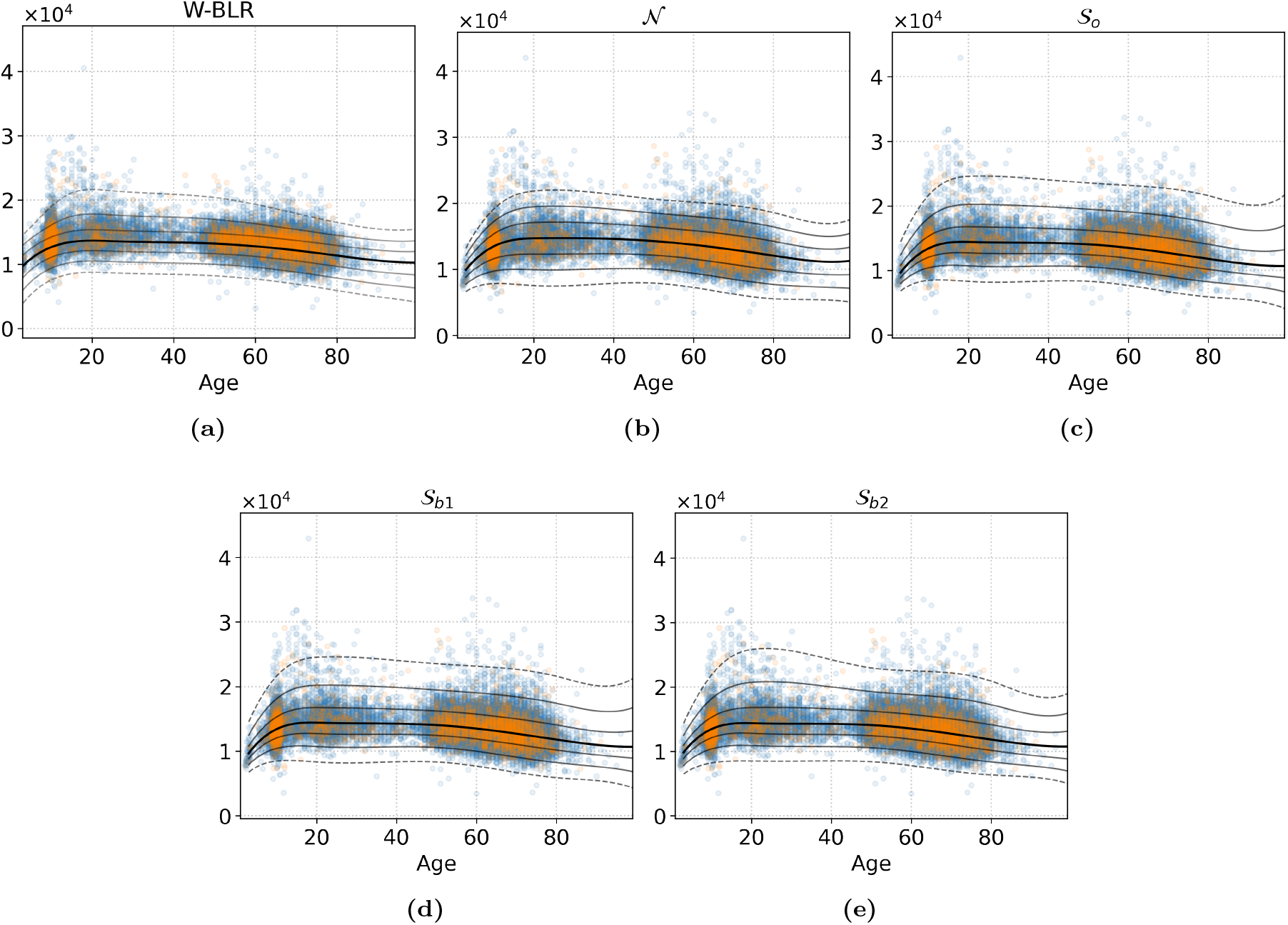
`Right-Cerebellum-White-Matter`

**Figure I.23:**
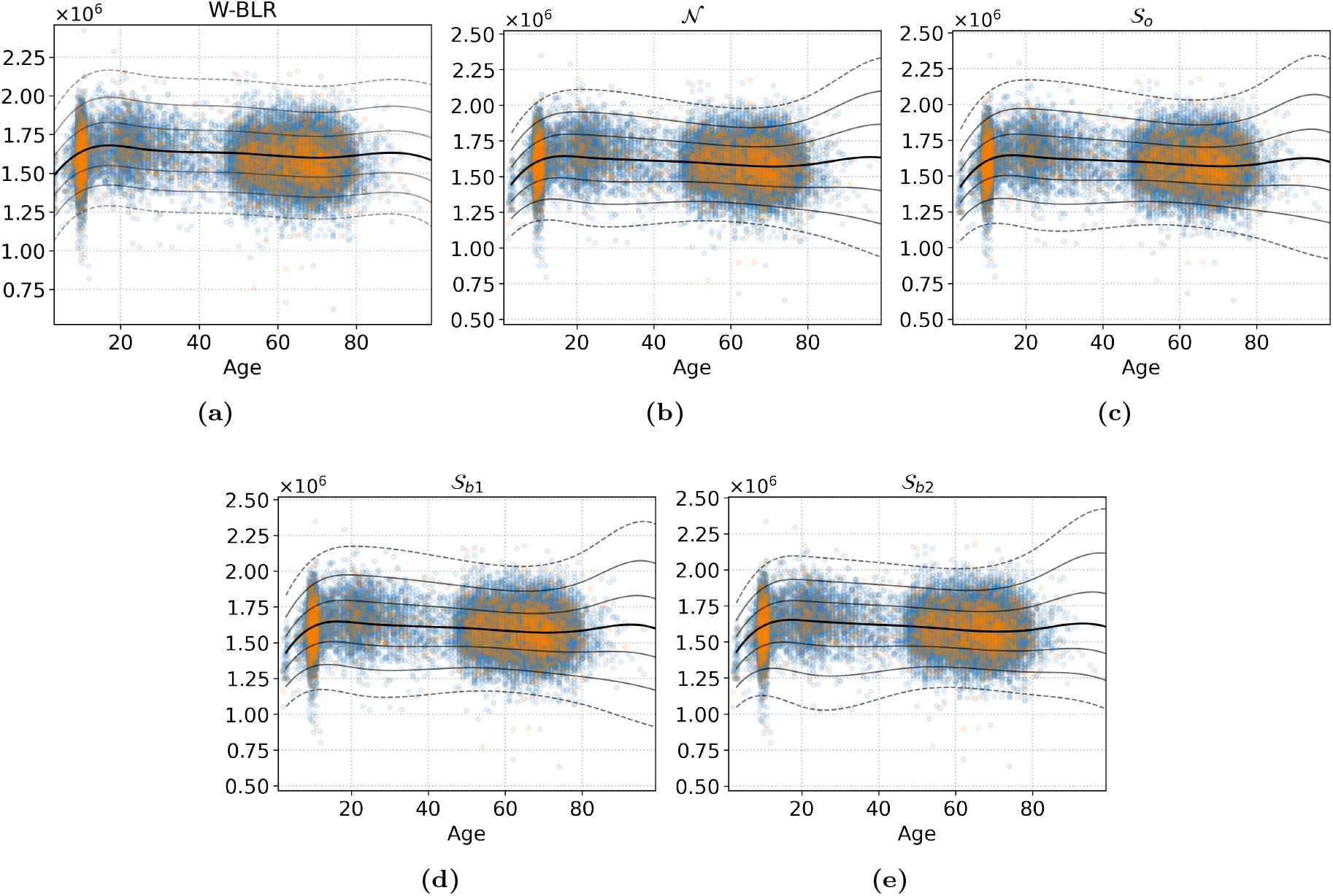
`EstimatedTotalIntraCranialVol`

**Figure I.24:**
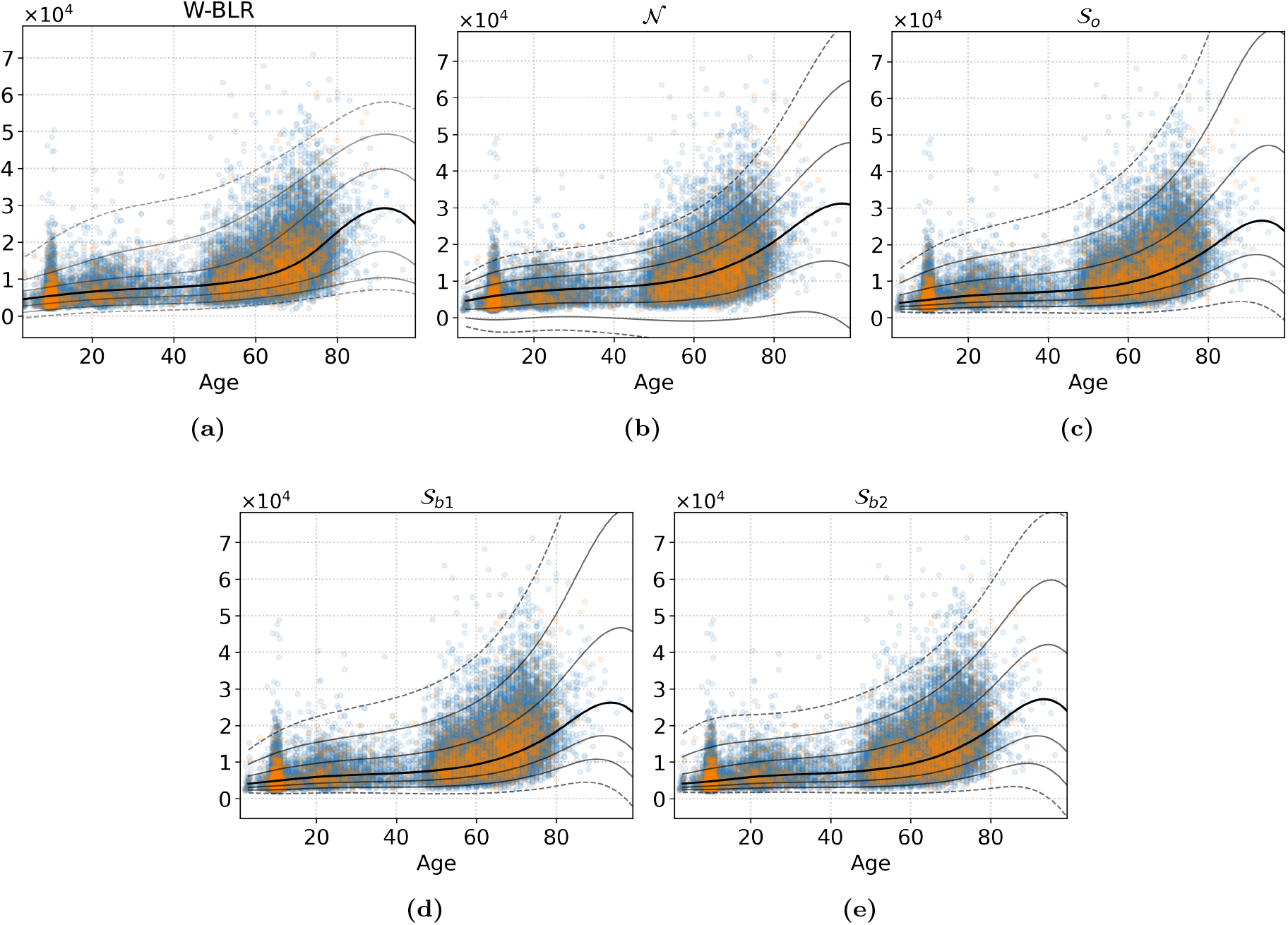
Right-Lateral-Ventricle

**Figure I.25:**
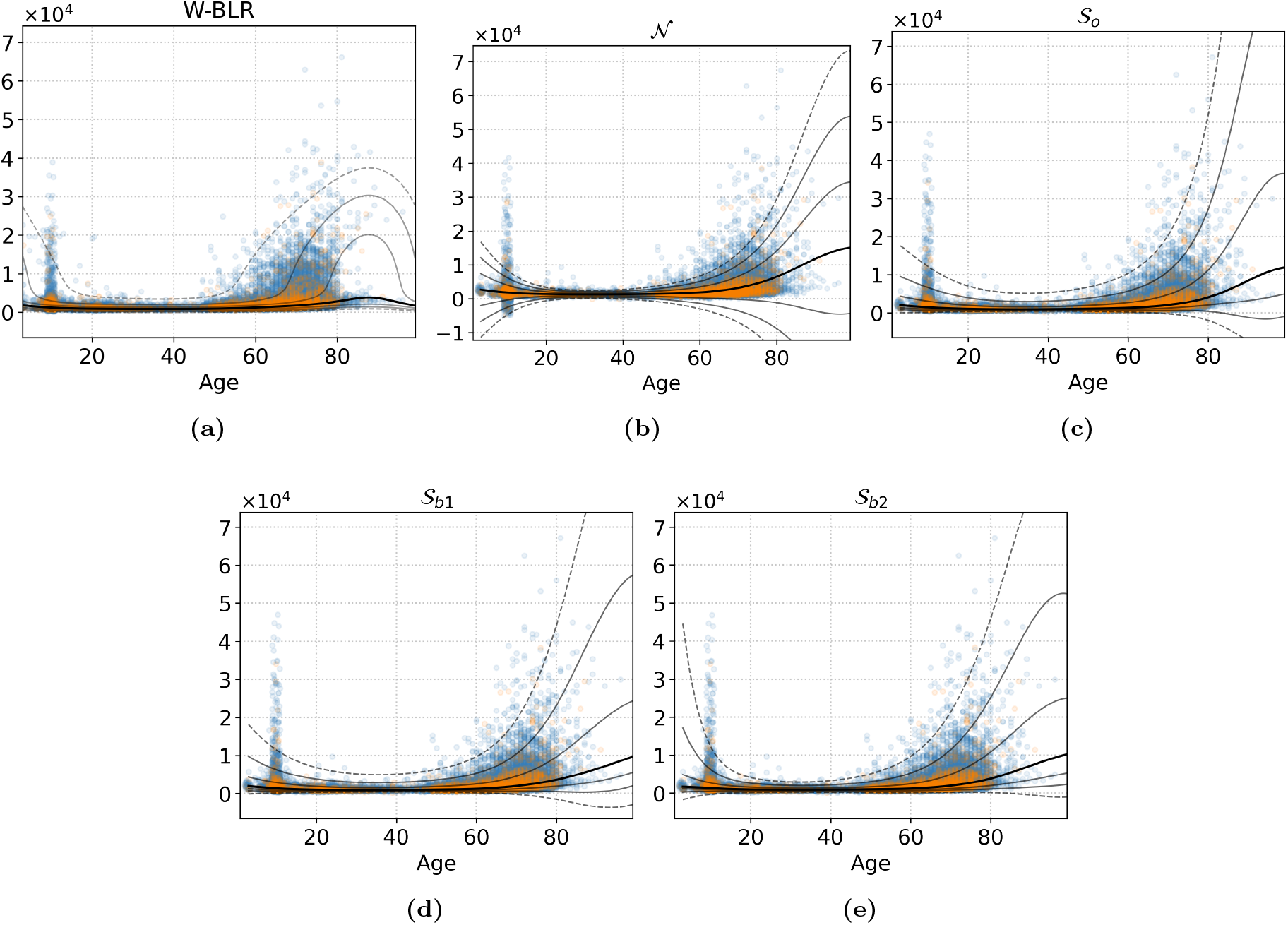
WM-hypointensities

**Figure I.26:**
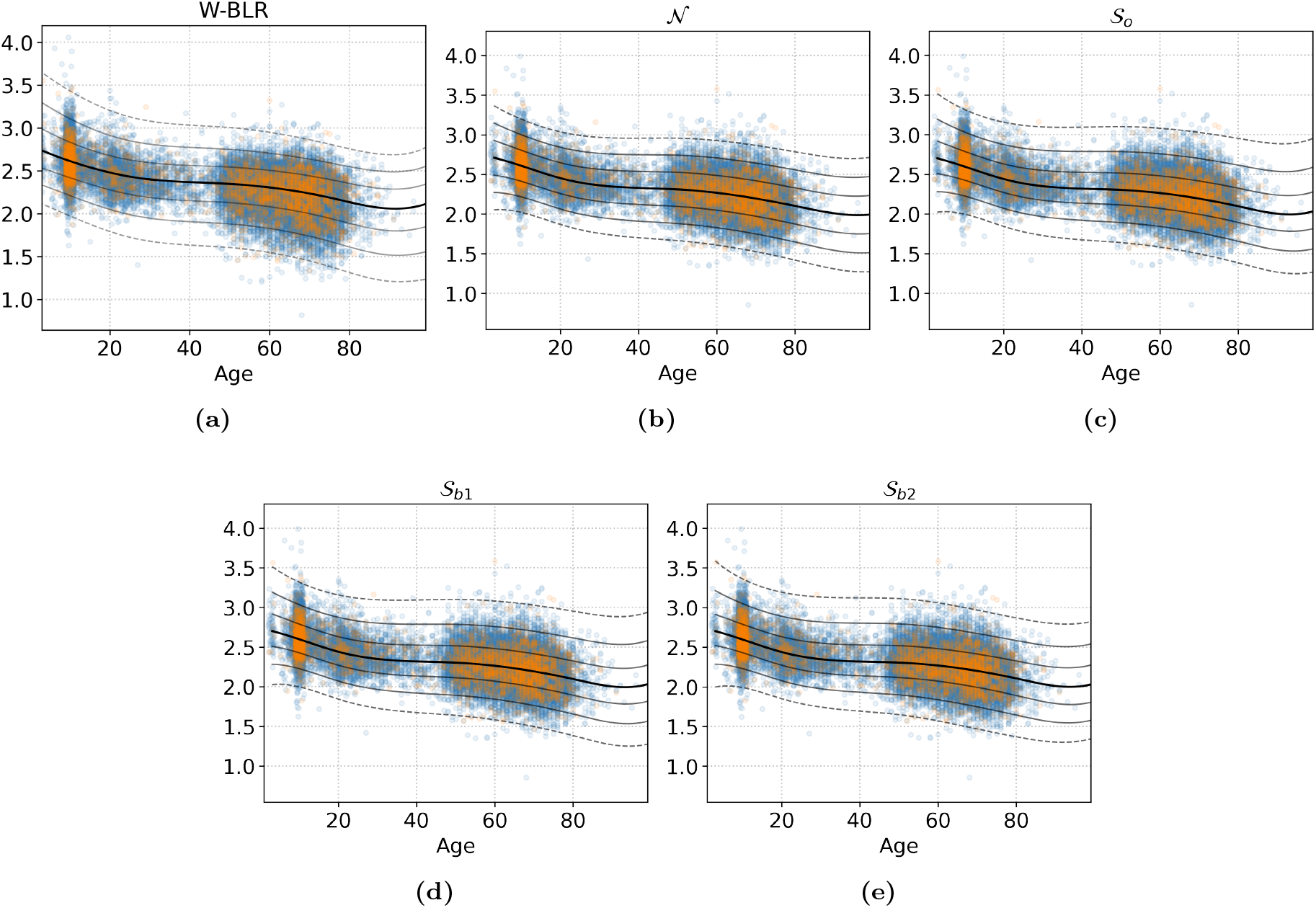
rh S interm prim-Jensen thickness

**Figure I.27:**
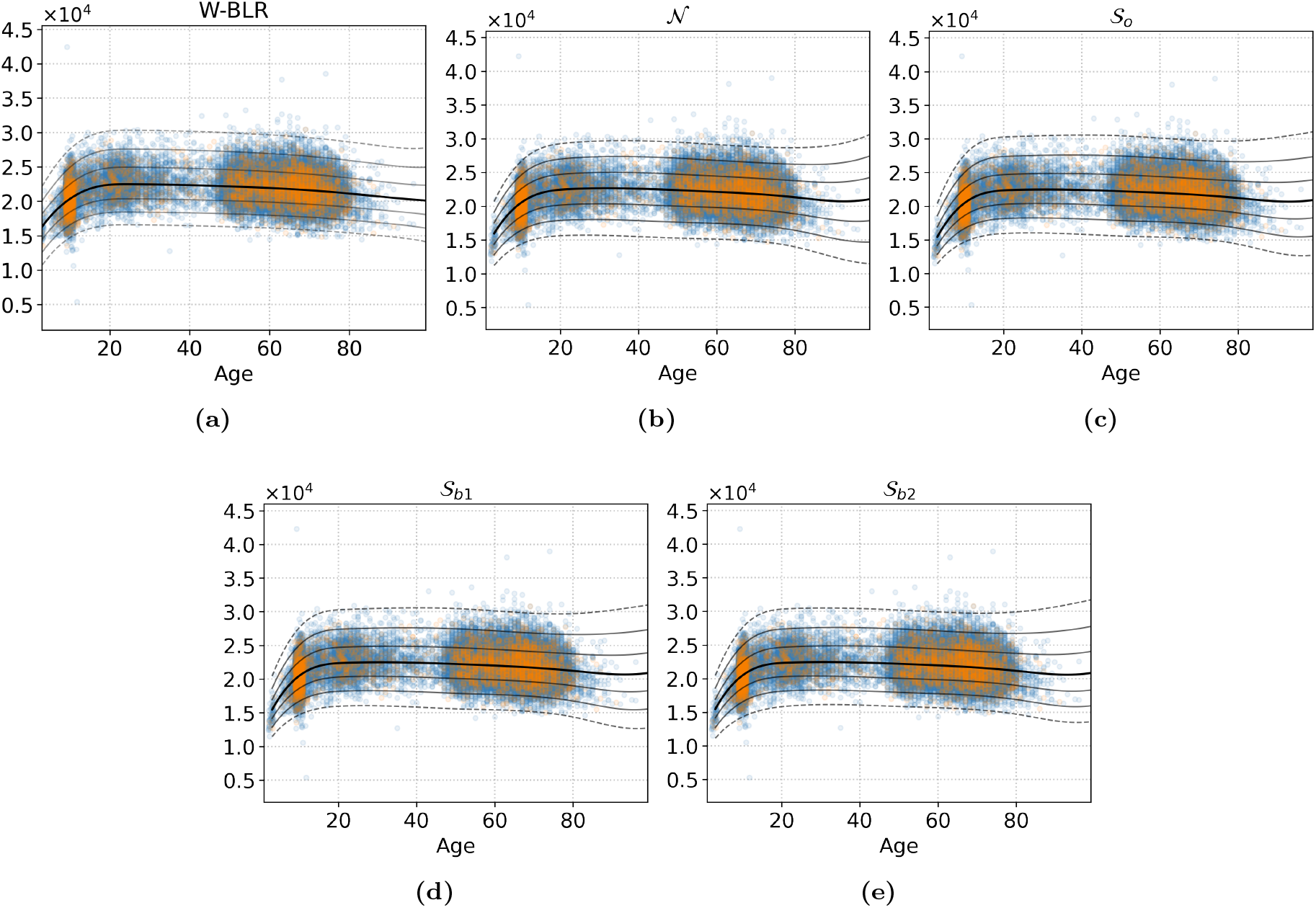
Brain-Stem

**Figure I.28:**
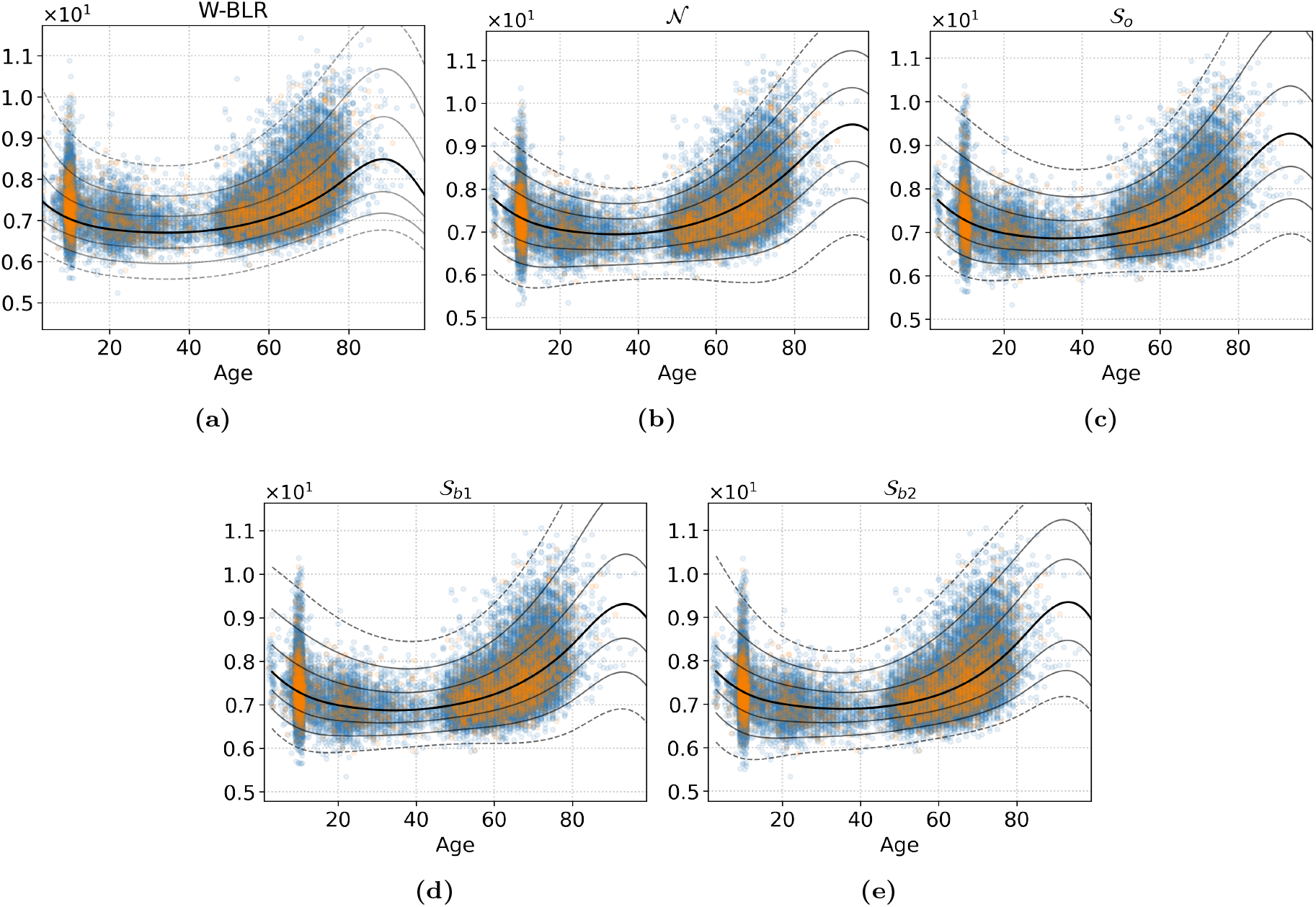
log WM-hypointensities

**Figure J.29:**
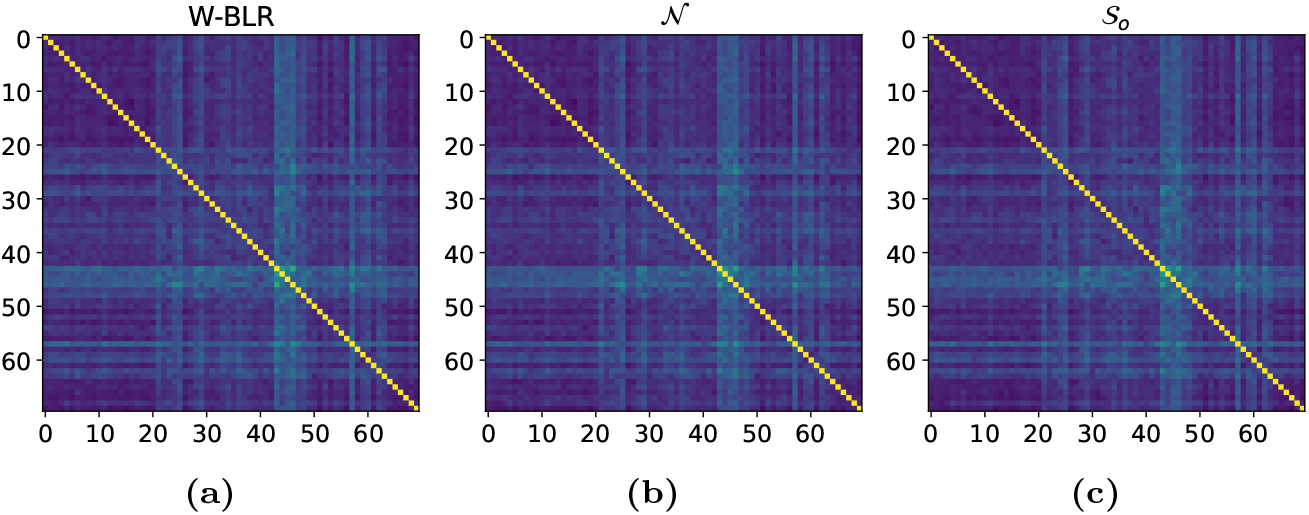
AUC scores of the Right-Cerebellum-White-Matter phenotype

**Figure J.30:**
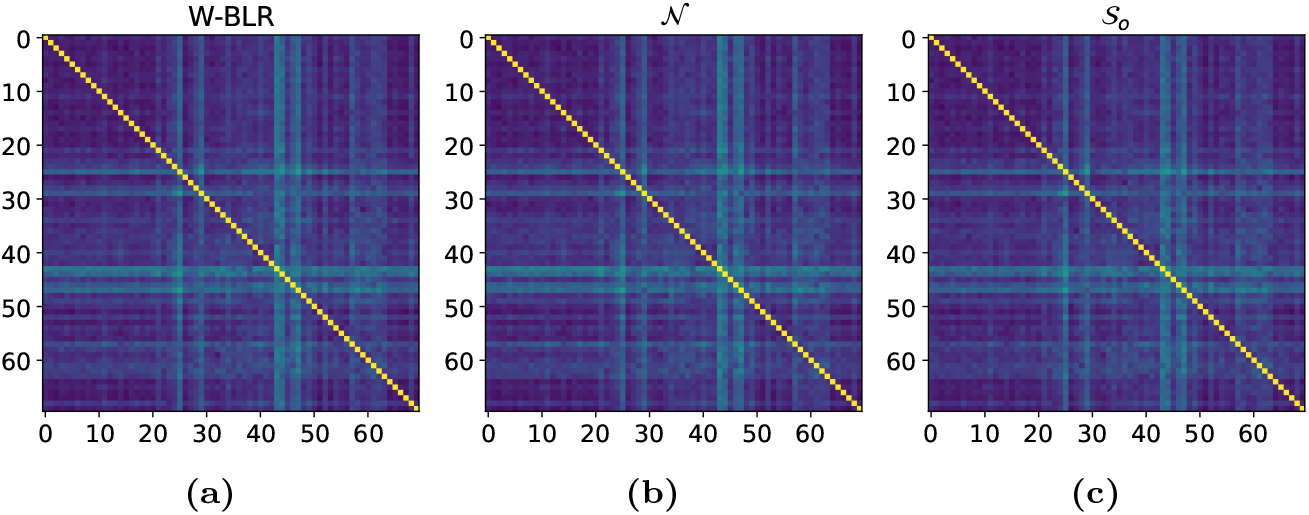
AUC scores of the Right-Lateral-Ventricle phenotype

**Figure J.31:**
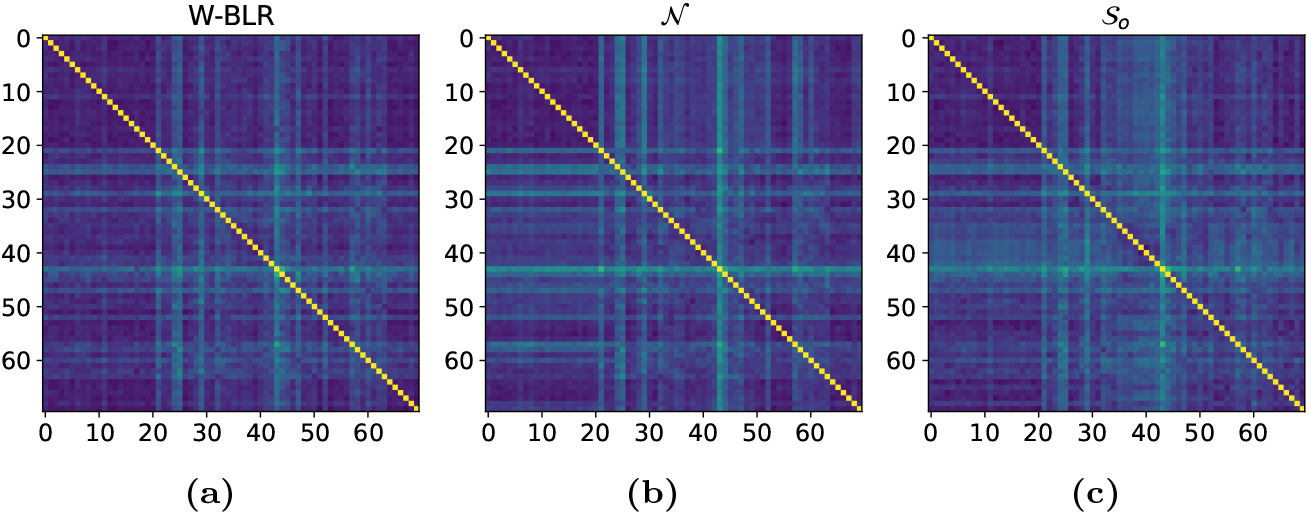
AUC scores of the WM-hypointensities phenotype

**Figure J.32:**
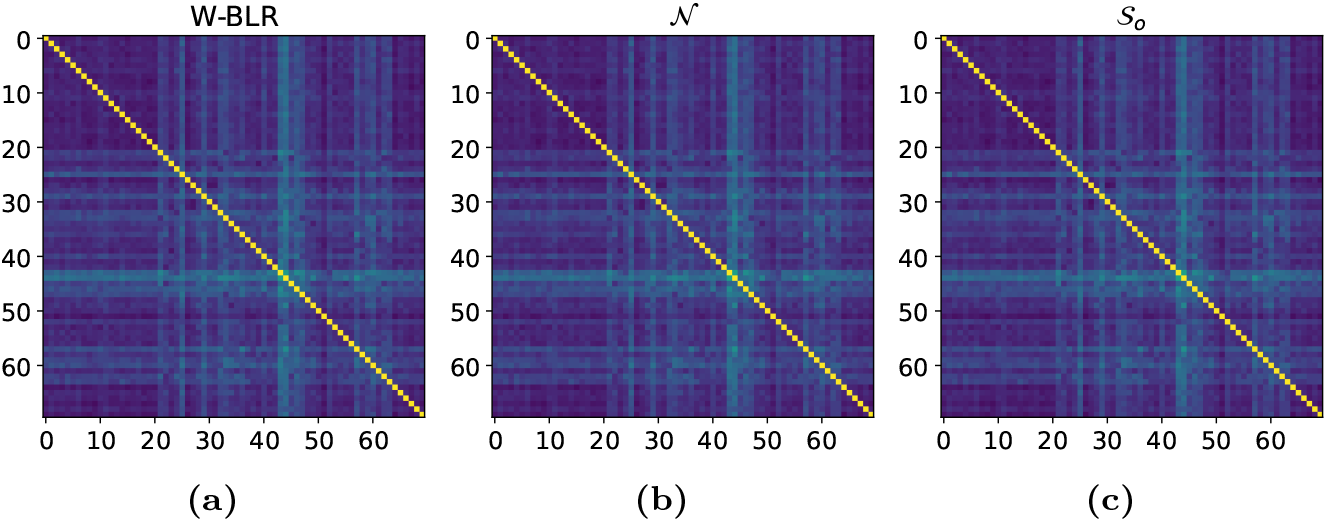
AUC scores of the EstimatedTotalIntraCranialVol phenotype

**Figure J.33:**
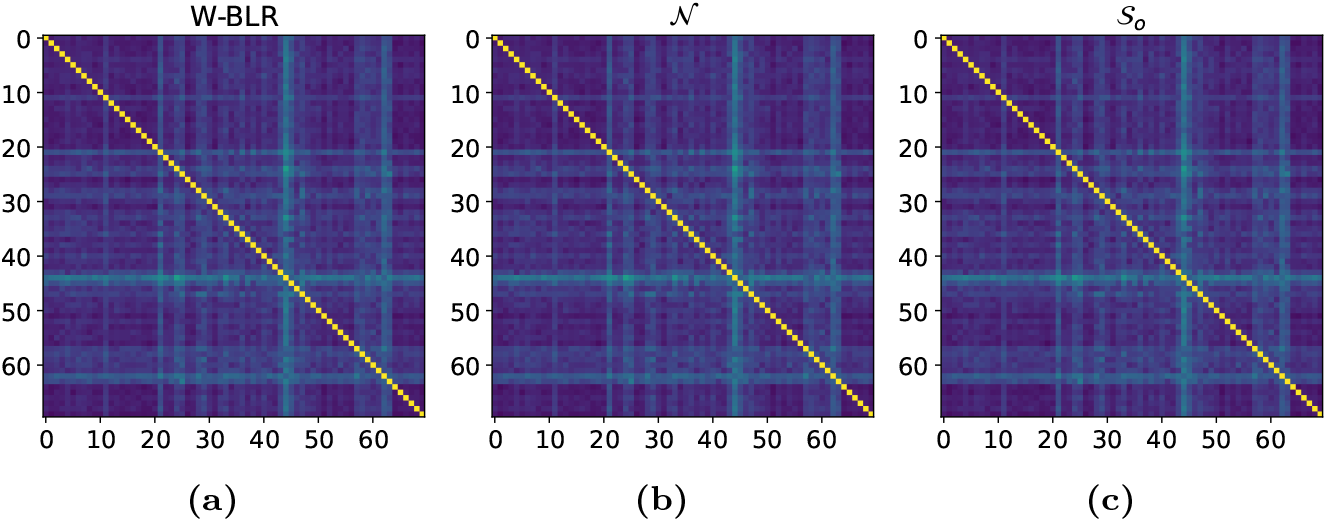
AUC scores of the rh S interm prim-Jensen thickness phenotype

**Figure J.34:**
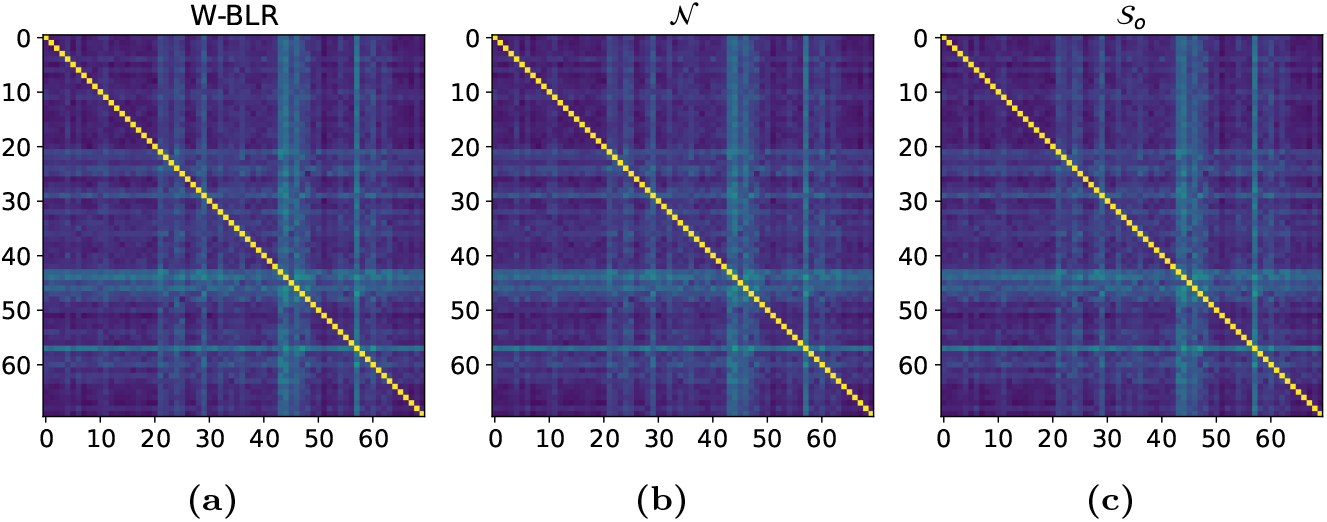
AUC scores of the Brain-Stem phenotype

**Figure J.35:**
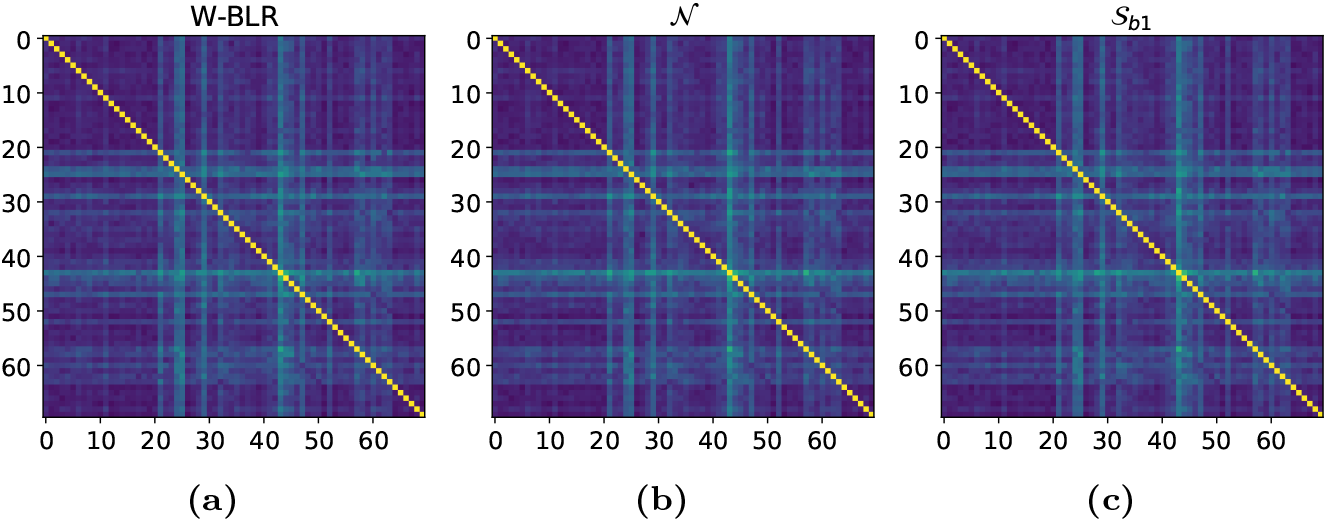
AUC scores of the Brain-Stem phenotype

**Figure J.36:**
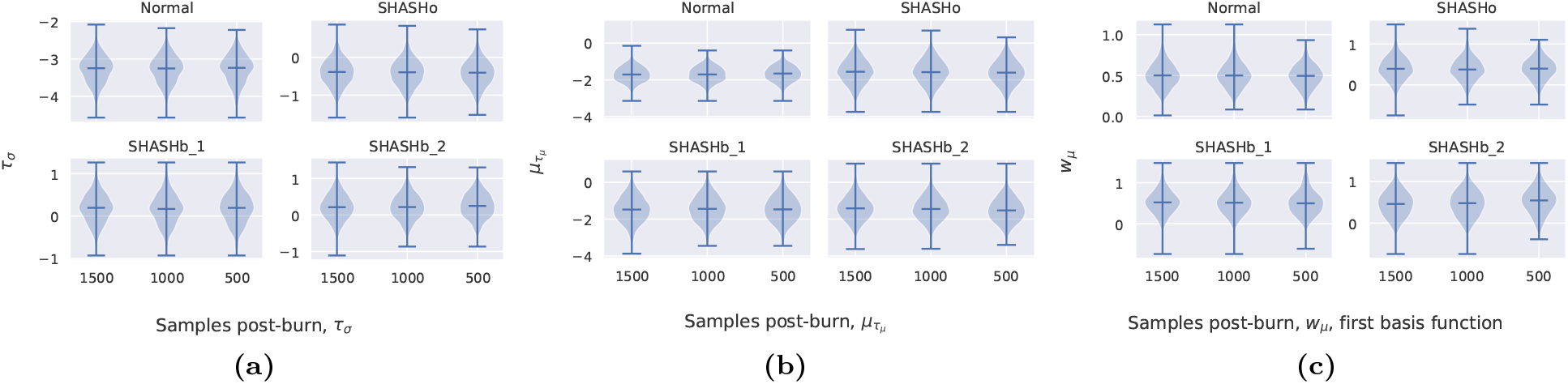
Effect of post-burn chain length on selected parameters.

##### Appendix B.1.1. Constant θ with batch effects

Combinations of site, sex, and possible other factors define distinct groups called *batches*. Here we use the term *batch effects* to indicate possibly unknown differences between known and distinct batches present in the data. HBR can deal with these batch effects in a principled way by modelling batch-specific parameters as deviations from a learned group mean. This allows information from all batches to flow into the hyperpriors, which in turn influence the estimated individual parameters for each batch. The models ℳ_**2a**_ and ℳ_**2b**_ encode this.

##### Model ℳ_2a_

CThe next simplest case is model ℳ_2*a*_, in which *θ* does depend on *Z* but not on *X*. This encodes the belief that the parameter is constant for all *x*, but that it may be influenced by batch effects. We sample the batch effects (*θ*_*b*_) as Gaussian deviations from a group mean (*µ*_*θ*_) with a group variance 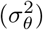, such that the sites can learn from each other. Additionally, we have that under the appropriate prior, the group variance acts as a regularizer.

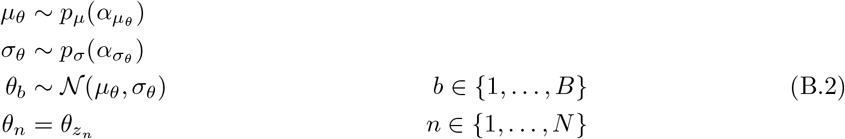

##### Model ℳ_2b_

The phenomenon known as the funnel is an issue that commonly afflicts hierarchical models, and it may occur in model ℳ_2*a*_. To give a brief explanation, when the parameter *σ*_*θ*_ is small, but we can not determine *θ* with enough confidence, the posterior distribution attains a wide support in the domain of *σ*_*θ*_, including very small values. If we sample in those regions where *σ*_*θ*_ is very small, the values *θ* all collapse to *µ*_*θ*_, and sampling of *θ* becomes very hard. Most samples of *θ* will be rejected, since the proposal distribution is tuned for a larger step size. For a more thorough explanation see [29]. In those situations where we fear a funnel may occur, we apply the following reparameterization as a remedy; the batch-specific deviation from the group mean is determined by sampling the group standard deviation (*σ*_*θ*_) from a positive distribution, and then multiply that with an *offset* (*ν*_*b*_) sampled from a central member of a symmetric distribution—like the standard Gaussian—for each *b* ∈ *{*1, …, *B}*. Sampling like this is sometimes called a non-centered sampling approach, as opposed to the centered approach in model ℳ_2*a*_. Here we propose model ℳ_2*b*_:

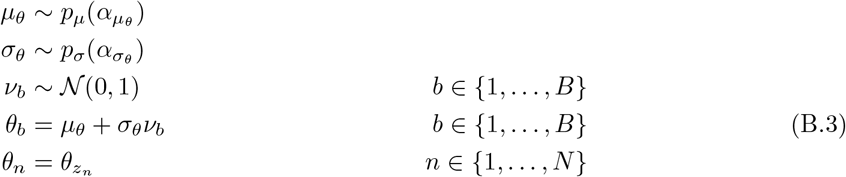

#### Appendix B.2. θ as a function of X

More complex models can be defined if we let *θ* follow a linear relationship with the basis expansion Φ(*X*) through a set of weights **w** ∈ **ℝ**^*dD*^ and an intercept *τ* ∈ ℝ. We sample **w** and *τ* from their own respective distributions, which is where the definitions from section Appendix B.1 are re-used.

##### Model ℳ_3.∗.∗_

Since **w** and *τ* are both constant parameters with possible batch effects, we can model them exactly like we did *θ* in ℳ_1_, or 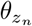 in models ℳ_2*a*_ and ℳ_2*b*_. The definitions there are general enough to allow modelling **w** as distributed according to a multidimensional density.

The subscript of these linear models consist of three terms, separated by periods; ℳ_3.∗.∗_. The first term is a 3, signifying that this is a linear model. The second and third terms indicate the model types that are used for **w** and *τ*, respectively. For instance: ℳ_3.1.1_ is a model where *θ* is a linear function of Φ through **w** and *τ*, which are both constants like *θ* in ℳ_1_. Model ℳ_3.2*a*.2*b*_ encodes for the model with a linear *θ*, where **w** and *τ* have unique values for each batch effect, but the former is modeled by the centered approach, and the latter by the non-centered approach explained on page 27. Both **w** and *τ* can thus be modeled in three distinct ways, allowing us to define a total of 9 different models with linear *θ* in this way. *θ* can thus be modeled in a total of 12 ways. 3 in which it is fixed, and 9 in which it has a linear dependence on Φ.

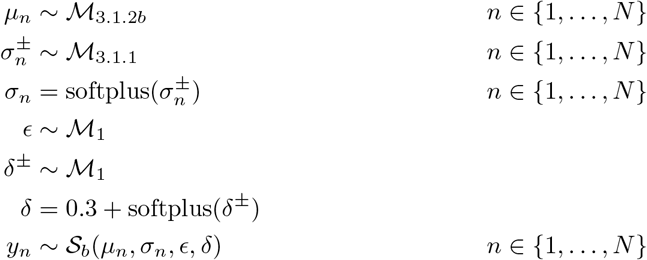

#### Appendix B.3. Hypothetical application

The proposed framework allows us to concisely specify how parameters are modeled.

##### Scenario

A pattern that is commonly observed in neurological phenotypes is the following: As the age of the participant (*X*) increases, the phenotype (*Y*) increases first (until around 25), and then starts to decrease. However, the variance in the phenotype increases with age as well. The data shows some positive skew, and may also be slightly kurtotic. Furthermore, the data is collected from various sites, so site-effects (*Z*_1_) are expected, and sex-related differences (*Z*_2_) need to be accounted for as well.

##### Modelling

Clearly, the parameter that controls the mean should be a function of age, but the non-monotonic behavior of the mean guides us away from a directly linear approach. Rather, we would use a polynomial basis expansion, or a regression spline [24]. We usually prefer the regression spline basis over a simple polynomial basis, because it does not induce global curvature [8]. A Gaussian distribution would not be able to capture the observed skew, so the decision for a likelihood function falls on the SHASH_*b*_, parameterized by *µ, σ, E*, and *δ*. Model ℳ_3_ is a fitting choice for both *µ* and *σ*, as they appear to be related with age. For *µ*, we expect that the batch effects do not affect the trajectory of the phenotype, but mainly influence the magnitude. Therefore, the weights **w** do not need to be modeled with batch effects, but *τ* does. The expected batch effects are not very large, so the non-central modelling approach (page 27) is a good choice. Model ℳ_3.1.2*b*_ is decided for *µ*. We do not expect batch effects in *σ*, but it does clearly have a relation with age. We can safely choose model *M*_3.1.1_ for *σ*. The last two parameters, *E* and *δ*, are both assumed to be constant across sites and age. So they are both modeled by ℳ_1_. To enforce positivity of both *σ* and *δ*, we apply a softplus transformation to the outputs of the corresponding models, and to fulfill the constraint discussed in section 2.1.2, we add a constant to *δ*.

We now have:

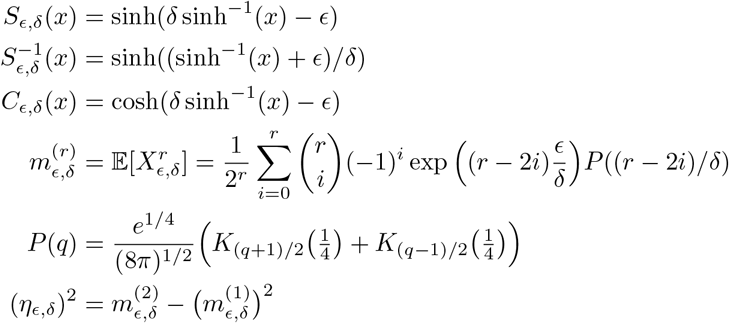

## Appendix C. Generative models

In all the generative models in this section, *D* is the dimensionality of the basis expansion *φ*. In our models this is taken to be 7. We repeat some earlier equations for quick reference.

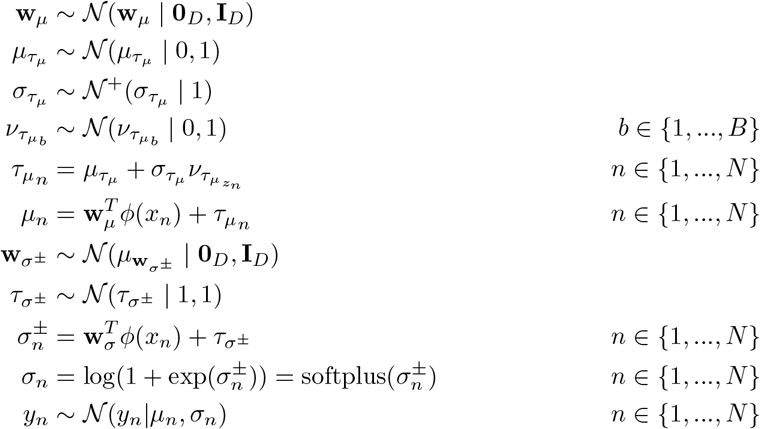

Where *K* is a modified Bessel function of the second kind.

### Appendix C.1. Model 𝒩

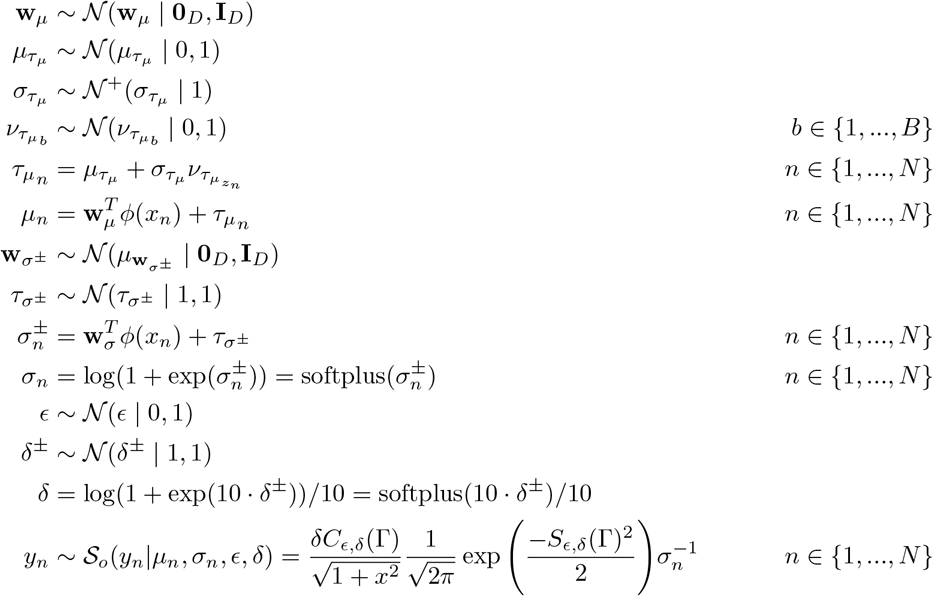

#### Appendix C.2. Model 𝒮_o_

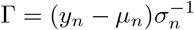

Where

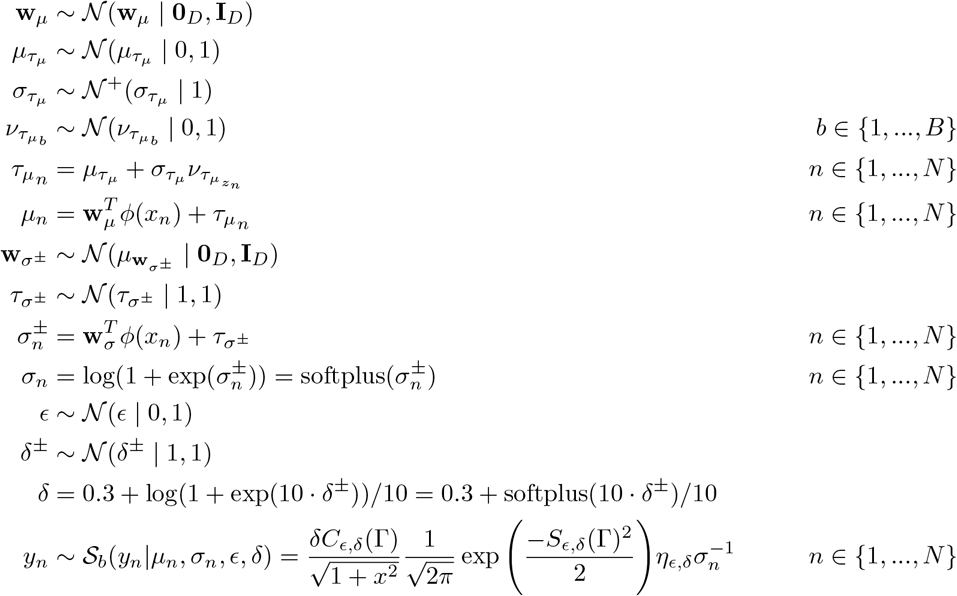

#### Appendix C.3. Model 𝒮_b1_

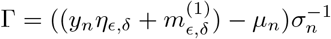

where

#### Appendix C.4. Model 𝒮_b2_

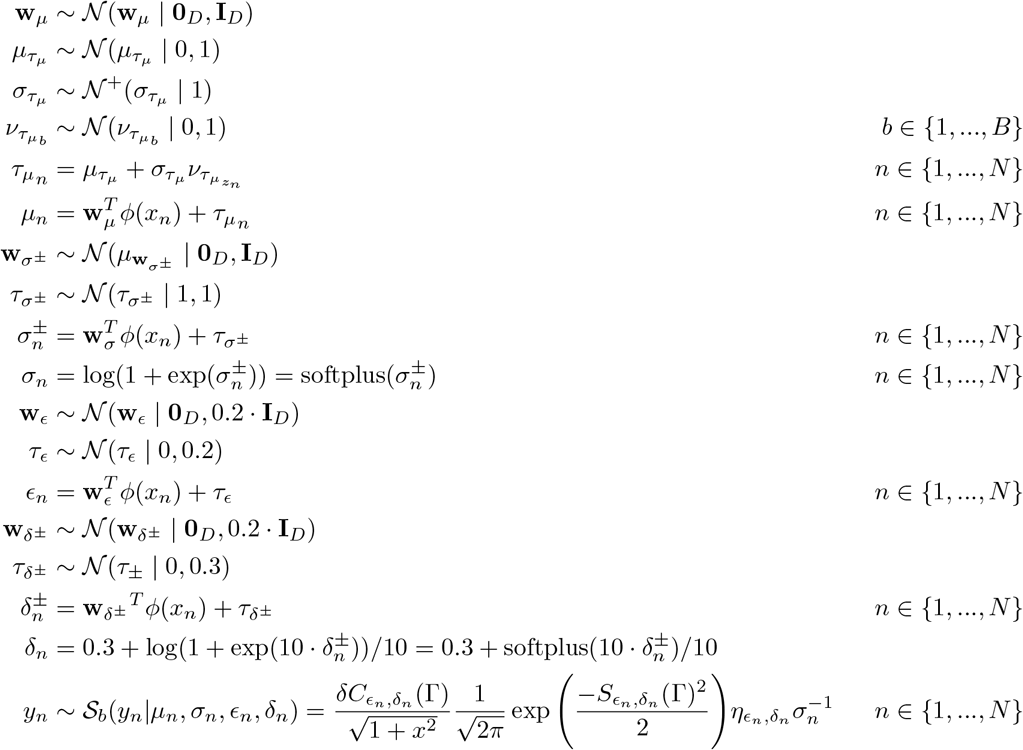

Where

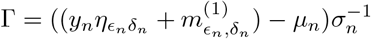

## Appendix D. MCMC sampling in a nutshell

Uncertainty quantification ideally accounts for all uncertainty in the model parameters, but is often hard to quantify exactly. MCMC approximates this uncertainty by sampling from the desired distribution. Given a generative model like above, Bayes’ rule gives us an expression for the posterior over our model parameters:

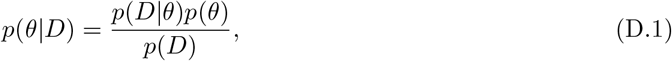

which we then use to compute the expectation of a function *f*:

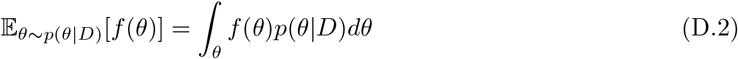

For instance, in normative modelling, *f* could return the deviation score of some data point. The expectation above gives us an estimate in which all the uncertainty in the model parameters is captured. Unfortunately, the integral in Eq. D.2 is intractable, and the denominator in Eq. D.1, *p*(*D*), contains an intractable integral as well:

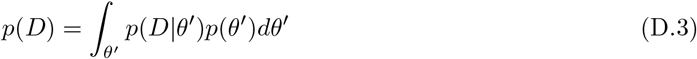

In practice it is impossible to determine *p*(*θ*|*D*) exactly, unless the prior is conjugate to the likelihood. MCMC samplers use a number of different methods to retrieve pseudo-random samples from *p*(*θ*|*D*), without ever needing to analyze the integral in Eq. D.3. With enough of these samples in hand, we use the following result:

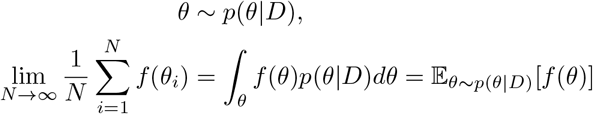

to approximate our desired expectation.

## Appendix E. Maximum-a-posteriori (MAP) approximation

Instead of doing an MCMC estimation, which is quite computationally expensive, another method is to find the MAP, and to compute the desired function based on that point.

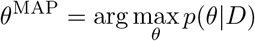

So in other words, the MAP estimate is just that point in parameter space that maximizes the posterior. It can be found using gradient based methods. Using the MAP-estimate is not the most principled method, because it does not account for the uncertainty in the parameters. For instance, the MAP estimate will always return a single *θ*, although the posterior could by multimodal, or contain large flat optima, which would justify multiple valid *θ*’s. However, the computational gains may be worth the loss of a representation of uncertainty.

## Appendix F. Data counts

## Appendix G. Empirical moments of SHASHo

## Appendix H. Empirical moments of SHASHb

## Appendix I. Percentiles of all features, by all HBR models

## Appendix J. AUC scores on all phenotypes

## Footnotes

1 The transformation of the Gaussian samples Z via the sinh-arcsinh transform only relates to the construction of the SHASH distribution, and it does not relate to z-scoring the data prior to analysis. Nevetheless we did standardize all variables used in the analysis before fitting since it can improve the performance of the samplers.

2 A differentiable function with a known and differentiable inverse

3 We have only real inputs because *y* is real and the cosh and sinh^−1^ functions map real numbers to real numbers.

4 Correlations like this cause problems for inference in HBR because of its reliance on MCMC sampling. In MCMC sampling, an algorithm traverses a path of pseudo-random points in parameter space, storing visited points as samples. Under the conditions that (i) the Markov chain has the posterior distribution as its stationary distribution, and (ii) detailed balance is satisfied, the samples that are retrieved follow the posterior distribution over model parameters, and can thus be used to approximate distributions. Correlations in the parameter space inhibit the sampler from efficiently exploring the parameter space, resulting in samples that do not follow the posterior very well, or do not converge to the target density at all.

5 Note that this differs slightly from the sample size in Rutherford et al. because we removed several small clinical samples for which permission for sharing was not available.

6 A one-hot encoding of an N-ary class label is a vector with a single ‘1’ at the index of the class that it labels, and zeros everywhere else.

## Notes

### Competing Interest Statement

CFB is a director and shareholder of SBGNeuro

### Summary of Updates

Additional analyses and clarifications following peer review

